# Protein docking on DNA surfaces using *de novo* designed binders

**DOI:** 10.64898/2026.11.25.762404

**Authors:** Raghu Pradeep Narayanan, Erika Snyder, Anh Pham, Ojas Gupta, Sruthi Raguveer, Konlin Shen, Cameron J. Glasscock, Meghan A. Morrissey, Shawn M. Douglas

**Affiliations:** Dept. of Cellular and Molecular Pharmacology, UCSF, San Francisco, CA 94143; Dept. of Molecular, Cellular, and Developmental Biology, UCSB, Santa Barbara, CA 93106; Dept. of BioSciences, Ken Kennedy Institute, Rice University, Houston, TX 77005; Dept. of Pharmaceutical Chemistry, UCSF, San Francisco, CA 94143

## Abstract

Positioning proteins on DNA-functionalized surfaces offers a way to organize catalytic, recognition and signaling activities across materials from synthetic particles to programmable nanostructures. However, site-specific attachment typically requires protein–oligonucleotide conjugation. Here we adapt a small, *de novo* designed protein that binds a six-base-pair sequence from one duplex face into a genetically encoded surface adapter. We define geometric constraints near blunt ends, hairpin loops and crossover junctions, and identify linker and site-spacing combinations that increase tandem *cis*-dimer affinity. In synthetic macrophage receptors, these dimers trigger phagocytosis of DNA-coated beads, preserve sensitivity to origami ligand spacing, and remain active after a 24-hour delay when DNA-conjugated receptors decline. We also design *trans*-dimers bridging neighboring origami helices to dock fluorescent proteins, enzymes, an antibody fragment and SpyCatcher cargo. Together, these design rules and binding modes establish a modular framework for genetically encoded protein–DNA interfacing that should generalize to new *de novo* binders.

---

DNA provides a programmable molecular interface for organizing interactions on material surfaces. DNA strands can be attached to nanoparticles, beads and other substrates, or assembled into nanostructures in which DNA itself forms the scaffold. These approaches combine sequence-specific recognition with control over the composition and geometry of the underlying material.^1^ Structural DNA nanotechnology^2^ produces scaffolds with defined size,^3–10^ shape^11–18^ and addressable positions.^19–24^ Such structures have been used to study viral entry,^25,26^ cellular signaling,^27–29^ molecular transport,^30–34^ subcellular organization^35^ and single-molecule dynamics.^36^ Attaching proteins to DNA-presenting surfaces brings catalytic, recognition and signaling functions to these materials.^37–41^ Doing so requires a reliable way to connect a chosen protein to a specified DNA site.

A commonly used approach is to covalently attach an oligonucleotide to a protein and recruit the conjugate through hybridization to complementary DNA on the surface. Preparing these conjugates requires chemical or enzymatic modification followed by purification, and the attached oligonucleotide can affect protein solubility or interfere with activity. For DNA origami, proteins are generally added after folding to avoid exposure to elevated temperatures. Access to attachment sites can then be limited by neighboring helices or other proteins; longer DNA handles improve accessibility but introduce flexibility between the protein and scaffold. These preparation and placement constraints become more demanding when several different proteins must be attached at distinct sites, each requiring a separately prepared conjugate.

DNA-binding proteins offer a genetically encoded route to attaching protein cargo to DNA. Zinc-finger, relaxase and HUH-tag adapters have been used to recruit proteins to DNA nanostructures,^42–46^ while transcription activator-like effectors (TALEs) have been used to assemble hybrid DNA–protein structures.^39,47^ Computational protein design has recently expanded the available DNA-binding architectures, producing small proteins that recognize short DNA sequences through major-groove contacts on one face of the duplex.^48^ This compact, single-face binding mode is well suited to DNA origami, where neighboring helices restrict access to internal duplexes and direct docking requires an interface that fits the exposed face of the DNA without clashing with adjacent helices. At the same time, contacting one face of a short site buries less surface area than encircling the duplex and makes access dependent on helical rotation. Coupling two single-face binders offers a way to increase affinity through avidity, either along a single duplex (*cis*) for linear DNA ligands or across neighboring helices (*trans*) on a multi-helix nanostructure.

Here we examine DBP35opt, a recently designed DNA-binding protein^48^ (Fig. 1a, b), across DNA substrates ranging from free duplexes and crossover junctions to DNA origami and cell-surface ligands. After defining local positional constraints near blunt ends, hairpin loops and crossovers, we pair two copies in *cis* across combinations of peptide-linker length and site spacing on a single duplex (Fig. 1c) and apply these dimers as genetically encoded macrophage receptors (Fig. 1e). To dock proteins at internal positions on DNA origami—where parallel helices present short duplex segments at defined spacing and rotational phase—we computationally design rigid *trans*-dimers bridging adjacent helices (Fig. 1d) and use them to place diverse protein cargos on an origami plate (Fig. 1e).

**Figure 1.**
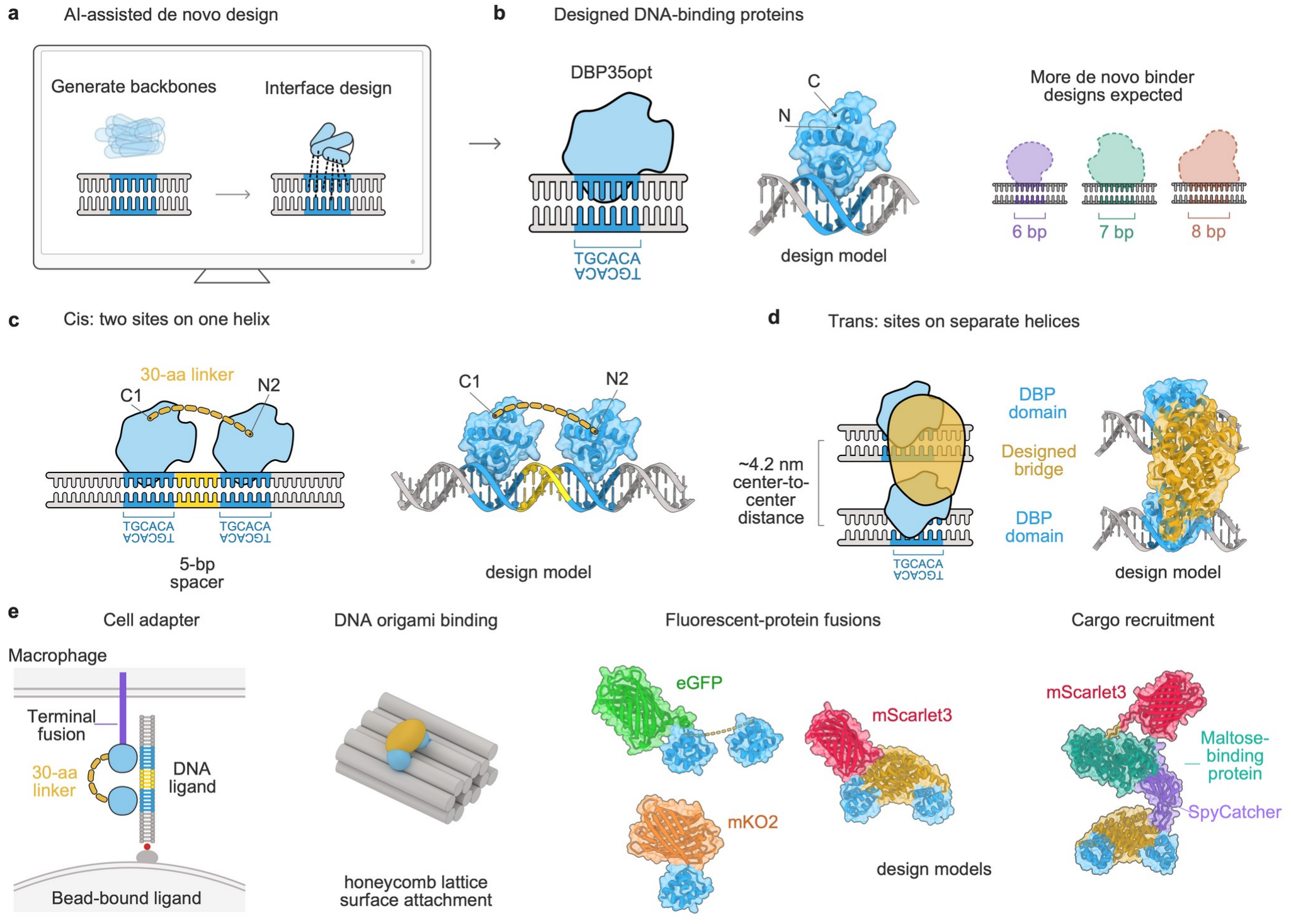
Designed DNA-binding proteins and fusion architectures. (a) Workflow for *de novo* backbone generation and DNA-interface design. (b) DBP35opt recognizes a six-base-pair target sequence (TGCACA) from one face of the duplex; dashed silhouettes illustrate potential binders with alternative footprints. (c) *Cis*-dimer architecture coupling two binding domains along one duplex via a flexible protein linker. (d) *Trans*-dimer architecture coupling two binding domains across separate helices via a designed protein bridge. (e) Fusion architectures for macrophage cell-surface receptors, DNA origami docking, fluorescent-protein labeling, and SpyCatcher–SpyTag cargo recruitment. Molecular structures are design models.

## Monomer binding rules on duplexes and junctions

We began with DBP35opt and DBP48, the two best-performing binders from a recent design campaign.^48^ We expressed and purified both and added them to pre-annealed hairpin duplexes (Methods). Only DBP35opt produced a clear shift (Supplementary Fig. S1a, b). We used DBP35opt for the subsequent experiments on target-site position and sequence. We walked the target site along a 13-bp duplex from the blunt end to the hairpin loop (Supplementary Fig. S1a) and tested a polyA duplex and a scrambled duplex as controls. The site had to lie at least two bases from the hairpin loop and two bases from the blunt end for DBP35opt to bind detectably, and no binding was detected on either control.

We appended DBP35opt to the C terminus of mKO2 and compared the fusion with the untagged binder. Native PAGE mobility-shift assays gave the same binding pattern as untagged DBP35opt (Supplementary Fig. S1e). By BLI, fusion slightly changed the dissociation constant from 17.4 nM to 19.7 nM, with a k_off_ of 6.27× 10^−4^ s^−1^ (Supplementary Fig. S5 and Supplementary Table S1). An mEGFP fusion behaved the same way (Supplementary Fig. S1c, d, f).

We next moved from a duplex to a branched structure, an immobile crossover junction (Fig. 2a; sequences in Supplementary Information). Varying the number of base pairs separating the six-base-pair recognition site from either the blunt end or the crossover, we observed binding whenever the site lay at least two base pairs from the blunt end and did not overlap the crossover (Fig. 2b, c). We then tested the binder on Q1, a custom asymmetric rectangular origami folded from a single-stranded scaffold produced in house (Supplementary Fig. S7a). Q1 presents 12 copies of the six-base-pair DBP35opt recognition site: six on terminal hairpins and six at internal positions along one face of the plate (one each on helices 0, 2, 4, 6, 8, and 10), shifted by one base pair from each site-bearing helix to the next to sample helical rotation. We incubated Q1 with the EGFP–DBP35opt fusion across a range of concentrations (Supplementary Fig. S7d). Negative-stain TEM showed some binding together with aggregation, and single-particle cryo-EM at fivefold molar excess gave no reliable density at the target sites (Supplementary Fig. S7b, c).

**Figure 2.**
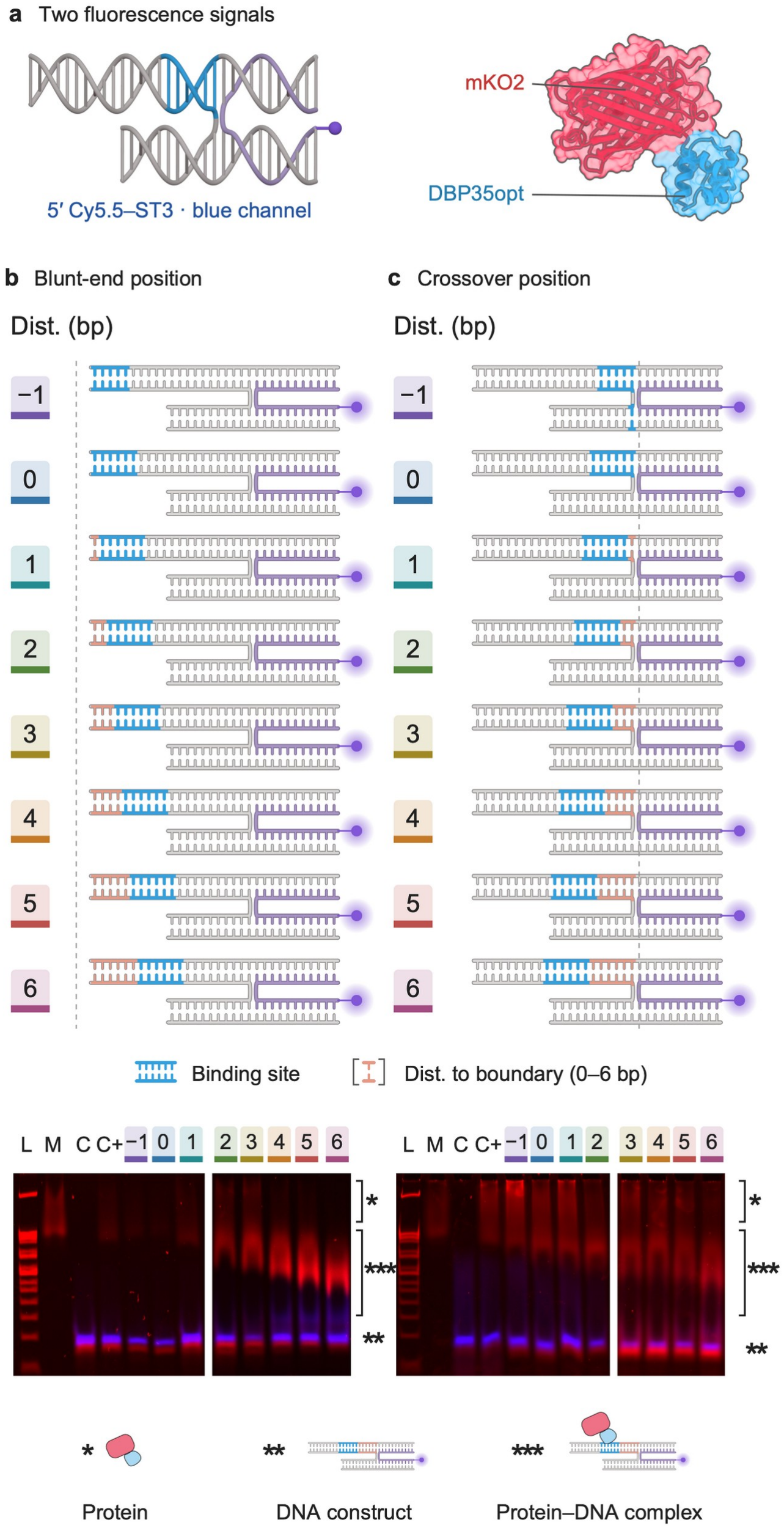
Local geometric constraints on monomer binding. (a) Schematic of the immobile crossover junction carrying a 5′ Cy5.5-labeled strand (ST3, blue channel) alongside a design model of the mKO2– DBP35opt fusion (red channel). (b, c) Recognition-site walks relative to a blunt end (b) or crossover junction (c), with corresponding two-color native PAGE gels below. Blue marks the six-base-pair recognition site and salmon marks its distance (0–6 bp) from the boundary; −1 places the site one base pair across the boundary (spanning five bases on the upper helix and one on the lower helix in c). Lanes: L, DNA ladder; M, protein alone; C, control junction lacking the target site; C+, control junction plus protein; −1 to 6, target junctions at the indicated boundary distances plus protein. White lines separate lanes from two gels run in parallel. For display in c, the red fluorescence channel of both gel segments was uniformly gamma-adjusted (γ = 0.55; Methods), while blue channels and b were unmodified. Single, double and triple asterisks indicate free protein, unbound DNA construct, and protein–DNA complex, respectively.

## Tandem *cis*-dimers across linkers and spacings

Because monovalent binding (∼20 nM *K*_D_) did not stably occupy Q1, we next asked whether coupling two copies of DBP35opt could increase affinity through avidity, starting with tandem *cis*-dimers that engage two sites along a single duplex (Fig. 3a). Protein linkers spanned 0 to 72 amino acids, and the spacing between the two target sites on the duplex spanned 0 to 12 bases. Across this matrix (Fig. 3b, c and Supplementary Figs. S2 and S3), the 12-, 18-, 27- and 30-amino-acid flexible-linker constructs all bound the 34-bp duplex containing a 5-bp spacer (34-5), while the no-linker *cis*-dimer bound the 2-bp-spacer duplex (34-2). The gel series resolved two shifted species whose relative mobilities correspond to one or two dimers bound per duplex (Fig. 3c). Heating the protein-duplex mixture to 55 °C for 10 min and returning it to 23 °C for 10 min shifted the band pattern from the species assigned to two dimers toward the species assigned to one dimer (Supplementary Figs. S2 and S3). We selected the 12- and 30-amino-acid flexible-linker constructs (12AAFL and 30AAFL) with the 34-5 duplex for BLI characterization and subsequent cell studies (sequences in Supplementary Information). BLI fits gave apparent dissociation constants and k_off_ values of 1.5 nM and 3.04× 10^−4^ s^−1^ for 12AAFL, and 0.124 nM and 4.25× 10^−5^ s^−1^ for the slow-dissociating component of a global two-component fit for 30AAFL (Supplementary Fig. S5, Supplementary Table S1, and Methods). Using 30AAFL, we varied the sequence flanking and separating the two target sites and tested the effect of a nick in the duplex; polyT spacers between the sites improved binding (Supplementary Fig. S4a–c). Excising the shifted band and re-running it returned a mixture of complex and free duplex (Supplementary Fig. S4d).

**Figure 3.**
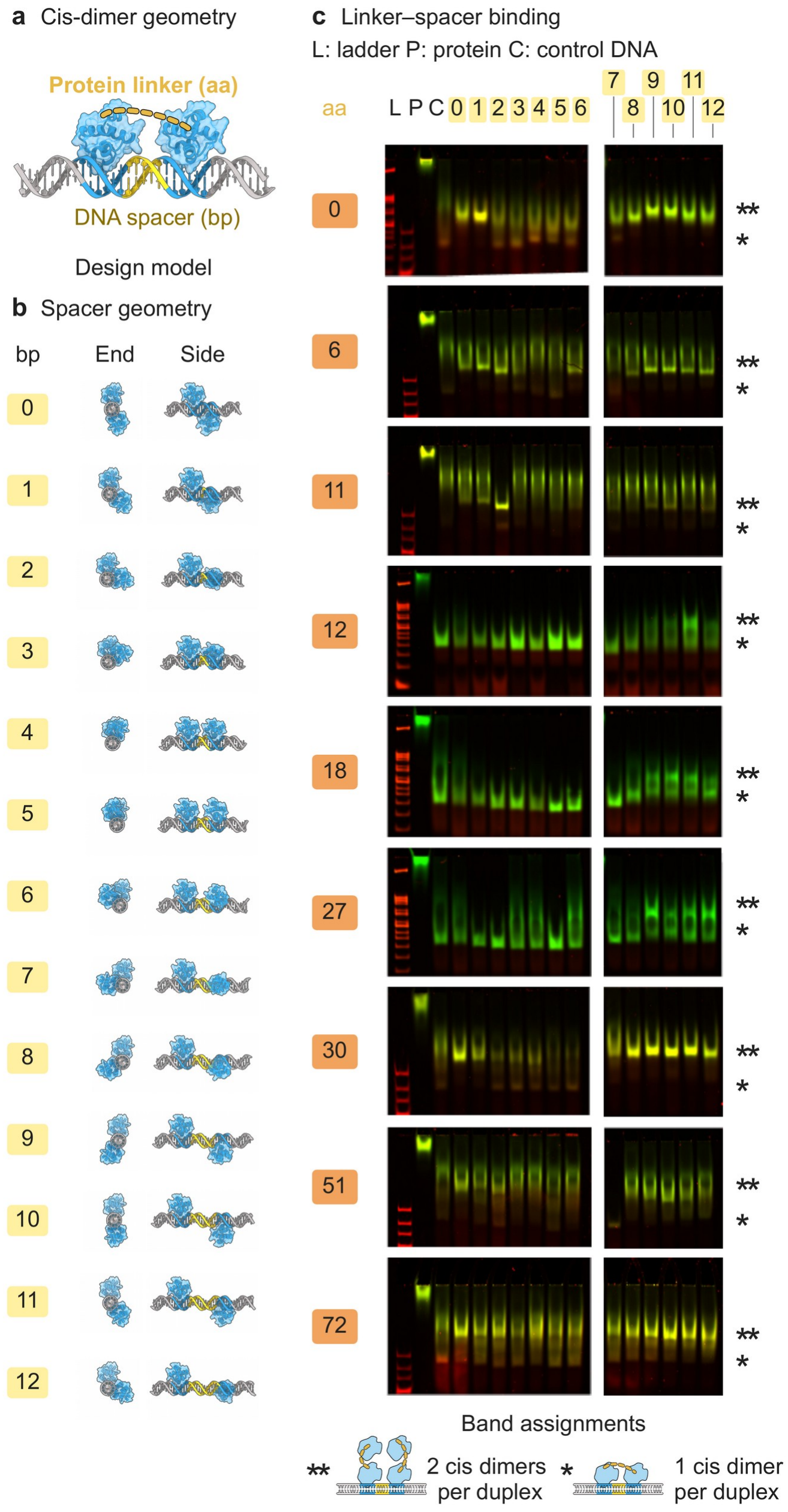
*Cis*-dimer binding across linker and spacer geometries. (a) Design model of a tandem *cis*-dimer bound to two sites on one duplex, defining protein-linker length (aa) and DNA-spacer length (bp). (b) End and side views of modeled duplexes across DNA spacers from 0 to 12 bp. (c) Native PAGE mobility-shift matrix spanning protein linkers from 0 to 72 aa and DNA spacers from 0 to 12 bp; the 6- and 11-aa constructs use rigid helical linkers (AEAAKA and AEAAAKEAAKA, respectively), whereas the 12–72-aa constructs use flexible (GGS)_n_ linkers. Lanes: L, DNA ladder; P, protein alone; C, control DNA lacking target sites. Single and double asterisks indicate shifted bands corresponding to one and two *cis*-dimers per duplex, respectively.

Single-particle cryo-EM of the no-linker and 30-amino-acid-linker *cis*-dimer complexes yielded 2D class averages and *ab initio* 3D volumes consistent with the overall dimensions of the duplex-bound dimers (Supplementary Fig. S6).

## Genetically encoded macrophage receptors

DNA nanotechnology allows precise patterning of ligands to test how their spatial organization controls receptor signaling in macrophages and T cells.^29,49^ This approach extends to native T-cell ligands: DNA origami displaying peptide–MHC complexes has revealed how ligand spacing, geometry and flexibility influence T-cell activation.^50^ Prior studies have used DNA chimeric antigen receptors (DNA CARs), in which an extracellular SNAP-tag is covalently conjugated to ssDNA and receptor activation is triggered by complementary DNA ligands on a target surface.^49,51^ We expanded on this approach by creating DNA-binding protein CARs (DBP CARs). The DBP CAR consists of an extracellular *cis*-dimer fused to the CD86 transmembrane domain and the intracellular signaling domain of the Fc receptor common gamma chain. The intracellular signaling domain of Fcγ receptors triggers phagocytosis, or engulfment, of foreign targets by macrophages when the extracellular domain is bound by its ligand. We expressed DBP CARs in RAW264.7 macrophages and provided them functionalized bead targets coated in DNA duplexes containing binding sites for the DBP CARs (Fig. 4a, b). DBP CAR macrophages phagocytose more bead targets with the addition of duplex DNA ligands. We next compared DBP CARs carrying 0-, 12- and 30-amino-acid linkers between the two ligand-binding domains (Fig. 4c). All three variants drove target-duplex phagocytosis, with the 12- and 30-amino-acid linkers reaching levels comparable to their IgG positive controls (adjusted P = 0.9946 and 0.1809, respectively; 0-amino-acid variant versus IgG, adjusted P = 0.0079) and all three linker architectures performing similarly to one another under target-duplex conditions.

**Figure 4.**
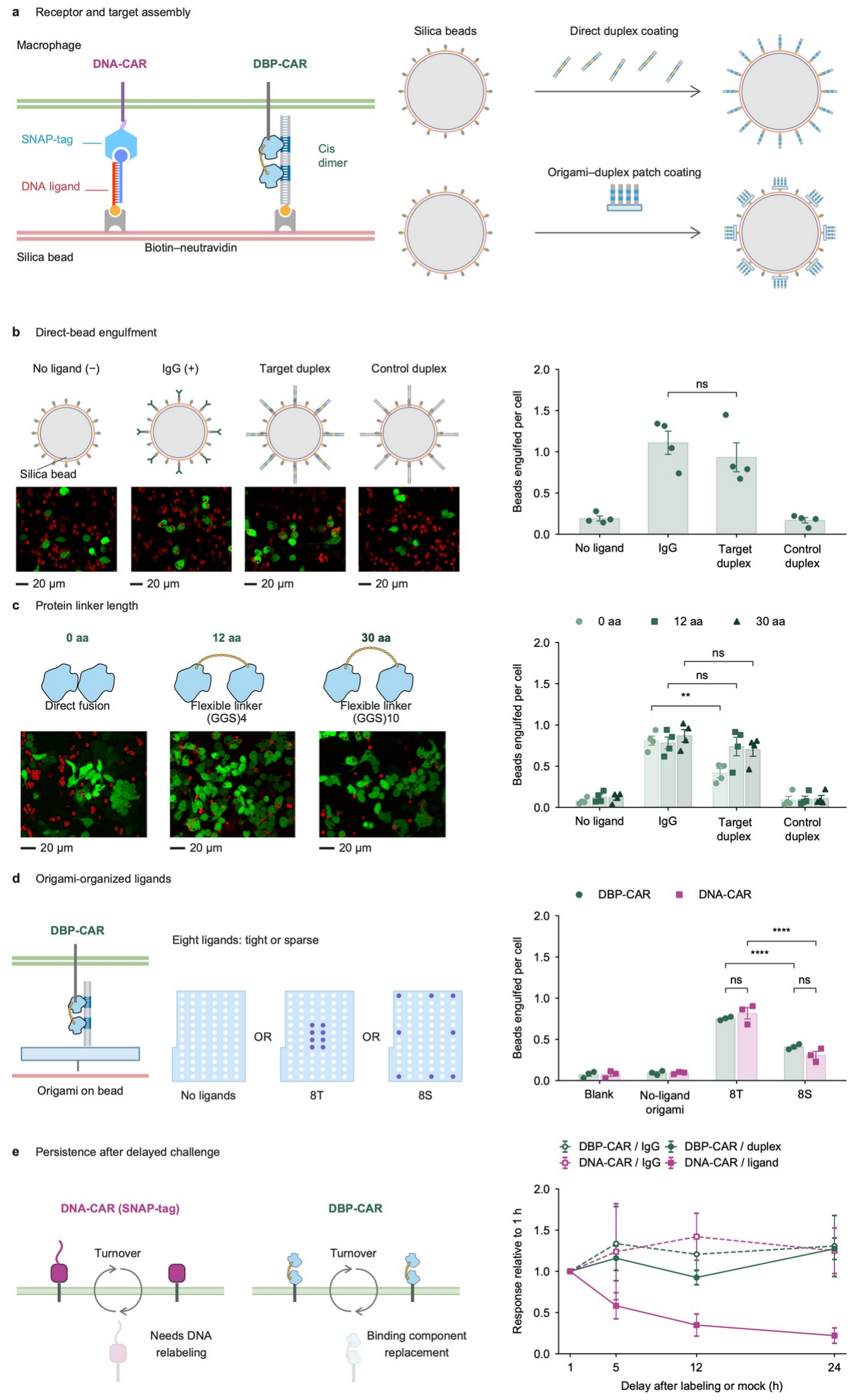
DNA-binding proteins as cell-surface adapters. (a) Receptor and target-bead assembly schemes for ssDNA-conjugated receptors (DNA-CAR) and genetically encoded *cis*-dimer receptors (DBP-CAR) engaging silica beads functionalized directly with DNA duplexes or with DNA origami pegboards. (b, c) Macrophage engulfment of duplex-coated beads across ligand controls (b) and DBP-CAR inter-domain linkers of 0, 12 and 30 aa (c). Confocal images show RAW264.7 macrophages (green) and silica beads (red); scale bars, 20 μm. (d) Engulfment of beads displaying origami pegboards without ligands or with eight tightly spaced (8T) or sparsely spaced (8S) ligands, alongside uncoated beads (Blank), for DBP-CAR and DNA-CAR macrophages. (e) Engulfment relative to the 1-h baseline when beads were added 1–24 h after ssDNA labeling (DNA-CAR) or mock treatment (DBP-CAR). Data show means ± SEM (n = 4 independent experiments in b, c, e; n = 3 in d), with individual experiment means overlaid in b–d. ns, not significant; **P < 0.01; ****P < 0.0001 (one-way ANOVA with Tukey test in b; two-way ANOVA with Tukey test in c and Šídák test in d).

Clustering of Fc receptors promotes signaling and phagocytosis.^49,52^ We varied the spacing of the DNA duplex ligands using DNA origami pegboards adapted from Kern et al.^49^ We created pegboards with tight (8T) and loose (8S) ligand spacing, conjugated them to supported-lipid-bilayer-coated bead targets, and incubated them with RAW264.7 macrophages expressing the medium-linker DBP CAR. Tightly spaced ligands triggered substantially more phagocytosis than loosely spaced ligands for both DBP CARs and DNA CARs (adjusted P < 0.0001 for each receptor; Fig. 4d), with DBP CARs matching the spatial-discrimination profile of DNA CARs on both the 8T and 8S designs (adjusted P = 0.9984 and 0.6966, respectively).

DNA CARs require a chemical conjugation step to attach their DNA recognition strand. DBP CARs encode the DNA-binding domain within the receptor itself. We compared phagocytic activity after delayed target addition: we challenged macrophages expressing either DNA CARs or DBP CARs with functionalized bead targets at increasing intervals after receptor ssDNA conjugation or mock treatment, respectively (Fig. 4e). DBP CARs retained phagocytic activity through the 24-hour time point, whereas DNA CAR activity declined relative to the 1-hour baseline.

## Designed *trans*-dimers for internal origami sites

While flexible *cis*-dimers are well suited to linear duplex ligands displayed on beads or pegboards, internal helices of a DNA origami lattice are interrupted by periodic crossovers and packed closely beside neighboring duplexes. Rather than placing two sites in tandem along a single helix, a multi-helix sheet allows two binding domains to bridge adjacent helices in *trans* through a rigid protein scaffold matched to their spacing and major-groove orientations (Fig. 1d). To test this architecture, we returned to the Q1 origami plate, whose internal target zone presents one six-base-pair DBP35opt site on each of six helices (0, 2, 4, 6, 8, and 10; Fig. 5a, b). Adjacent site-bearing helices form five candidate *trans*-dimer pairs—(0, 2), (2, 4), (4, 6), (6, 8), and (8, 10)—and advancing the target sequence by one base pair from each site-bearing helix to the next rotates the major groove progressively across the array (Fig. 5c). Both the two sites within any pair and successive pairs across the plate therefore differ by a single base pair of helical phase.

**Figure 5.**
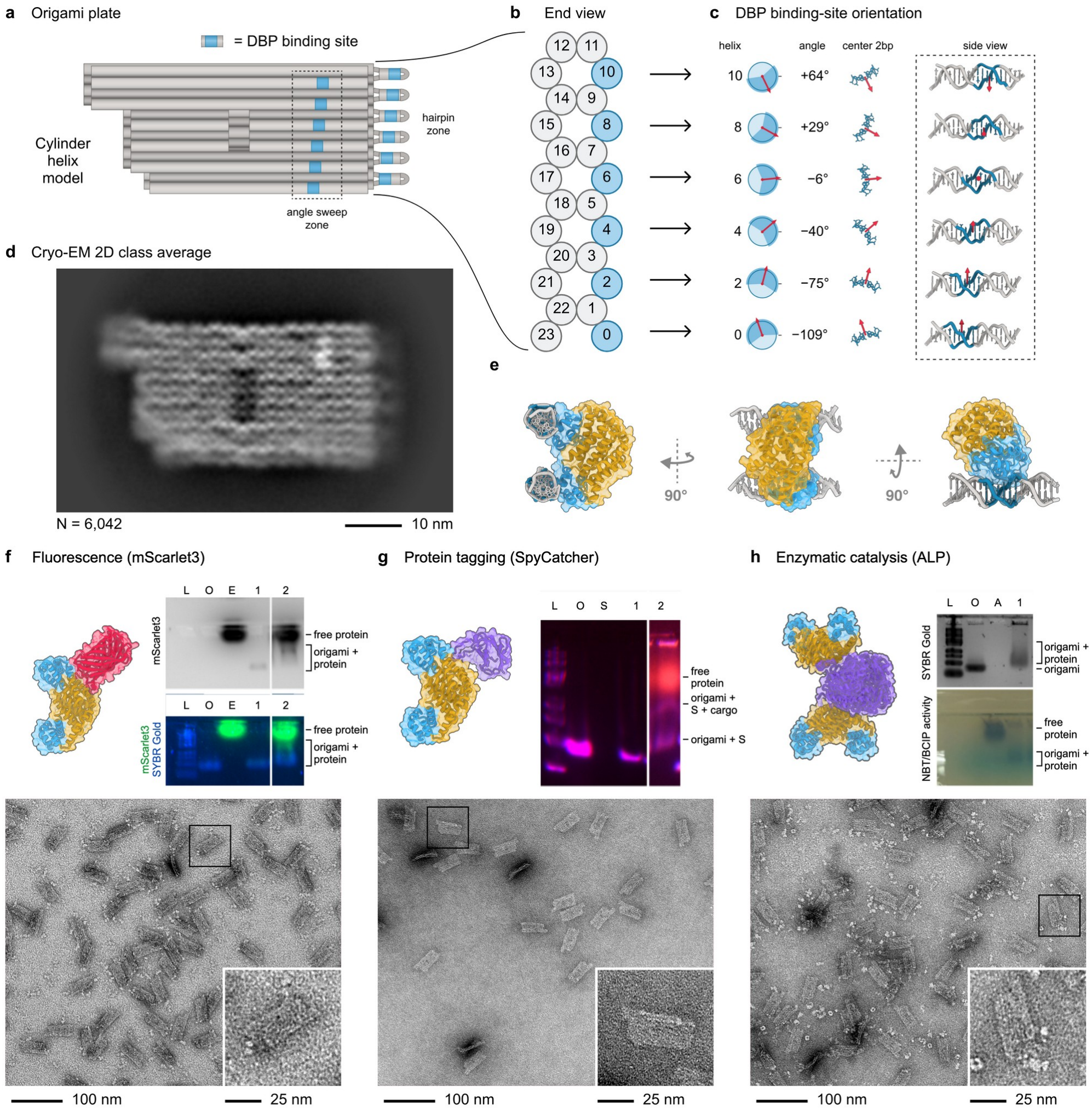
Geometric recognition and protein cargo recruitment on DNA origami. (a) Cylinder model of the Q1 DNA origami plate showing six internal DBP35opt recognition sites (angle-sweep zone) and six terminal hairpin sites in blue. (b) Cross-sectional end view numbering helices 0–23, with the six internal site-bearing helices (0, 2, 4, 6, 8, and 10) highlighted in blue. (c) Major-groove orientations across the six internal sites, which advance by one base pair per site-bearing helix across 173° (+64° to −109°; red arrows mark site-center directions and blue sectors show the six-base-pair groove sweep), alongside axial views of the central two base pairs and side views of each duplex segment. (d) Cryo-EM 2D class average of Q1 bound by the D4 *trans*-dimer (N = 6,042 particles), shown at the same scale as a; scale bar, 10 nm. (e) Design model of the D4 *trans*-dimer bound to the target duplexes of helices 8 and 10 in three views (∼90° rotations), with DBP35opt recognition domains in light blue, the designed bridge in gold, and six-base-pair DNA target sites in dark blue. (f–h) Design models, agarose gel assays, and negative-stain TEM micrographs for D4 fused to mScarlet3 (f), SpyCatcher3 recruiting SpyTag3–MBP–mScarlet3 cargo (g), and alkaline phosphatase (h). Lanes: L, 1-kb DNA ladder; O, Q1 origami alone; E, S and A, the respective D4 fusions alone; 1 and 2, Q1 with a sixfold (lane 1) or 48-fold (lane 2) molar excess of the fusion (f) or pre-assembled SpyCatcher–SpyTag cargo complex (g). White lines separate lanes juxtaposed from the same gel. Panel f shows mScarlet3 fluorescence before staining (top) and composite mScarlet3 (green) plus SYBR Gold (blue) after staining (bottom); panel g shows composite mScarlet3 (red) and SYBR Gold (magenta); panel h shows SYBR Gold staining (top) and NBT/BCIP phosphatase activity staining (bottom) of the same gel. Insets show magnified views of the boxed particles; scale bars, 100 nm (overviews) and 25 nm (insets).

Focusing on the helix 8–10 pair, whose major grooves face into the solvent-accessible cleft between the two duplexes (Fig. 5c), we designed *trans*-dimers *in silico* as single polypeptides coupling two DBP35opt recognition modules. One module was circularly permuted around the second using *de novo* connecting segments and terminal extensions so that two structured bridges fixed their relative geometry (Supplementary Fig. S8; full sequences in Supplementary Information). After placing two copies of DBP35opt onto the helix 8 and 10 target sites in PyMOL to capture their array spacing and orientation, we generated ∼1,000 fused protein backbones per condition with RFdiffusion (v1), designed sequences for the newly built regions with ProteinMPNN while holding the DNA-contacting residues fixed, and ranked candidates by standalone AlphaFold 3 fold self-consistency (Methods).

We expressed the four top-ranked designs—D4, D10, D34, and D46—and screened their binding to Q1 by negative-stain TEM. Based on manual counting of negative-stain TEM images, D4 gave the highest occupancy at the internal target zone, reaching 44% compared with 18% for the next-best design, D34 (Supplementary Fig. S10 and Supplementary Data 1), and we selected D4 for cryo-EM and cargo-attachment experiments. Single-particle cryo-EM 2D classification yielded a class average with clear additional density localized over the intended helix 8 –10 target region of the plate (Fig. 5d), consistent with the position and scale of the modeled two-helix complex (Fig. 5e).

## Modular docking of diverse protein cargos

We then used D4 to place protein cargo on Q1. We built fusions representing four classes: (i) the fluorescent protein mScarlet3, (ii) an anti-mEGFP antigen-binding fragment, (iii) the enzymes alkaline phosphatase and β-galactosidase, and (iv) the adaptor SpyCatcher, which recruits SpyTag-modified cargo. Negative-stain electron microscopy showed all four classes incorporated onto the origami surface (Fig. 5f–h and Supplementary Fig. S9). Gel assays further confirmed mScarlet3 fluorescence on Q1, SpyCatcher-mediated recruitment of SpyTag–MBP– mScarlet3 cargo, and colorimetric turnover by origami-bound alkaline phosphatase (Fig. 5f–h).

## Conclusions

For the DBP35opt-based constructs and assay conditions reported here, binding depended on the position and accessibility of the DNA recognition site. In the tested substrates, binding was reduced or undetectable when sites were too close to blunt ends or hairpin loops, or overlapped crossover junctions. Monomeric binding did not produce reliable density at the origami target sites, while linking two copies in *cis* increased affinity on duplex substrates in a manner dependent on linker length and site spacing. We also designed *trans*-linked dimers for paired sites on neighboring origami helices and used them to attach protein cargo. DBP35opt-based *cis*-dimers supported ligand-dependent phagocytosis when incorporated into receptors displayed by macrophages. Together, these results show how matching binder valency and linkage geometry to the local DNA architecture enables genetically encoded docking across both nanostructured scaffolds and living cells.

The position-dependent binding observed here is consistent with differences in access to the recognition site. In the origami model, shifting the site along a helix changes the orientation of the major groove relative to neighboring helices. End fraying or local changes in DNA conformation near crossovers could also contribute to the reduced binding at those positions. In the *cis*-dimer matrix, the joint dependence on protein-linker length and DNA-spacer length reflects the interplay of helical phasing, steric access and linker flexibility.

The *trans*-dimer architecture uses two connecting segments to couple the recognition modules. Designing both connections around a pair of DNA sites provides a way to specify their relative orientation. Similarly, the shift toward the single-dimer species after brief thermal annealing indicates that mild heating promotes equilibration toward bivalent engagement on a single duplex.

Because these complexes are noncovalent, they remain in dynamic equilibrium and can partially dissociate upon strong dilution or gel extraction, whereas applications requiring permanent covalent linkage can draw on genetically encoded ssDNA-conjugating tags.^45,46^ Bivalency is what makes a noncovalent interface practical here: the dimers bind tightly enough that irreversibility is not required for the assays we report. The tradeoff is in targeting flexibility, since a dimer needs two sites at a defined spacing and helical phase where a monomer needs only one. We established these rules with one parent binder. Follow up studies would demonstrate the rules extend to a second designed binder and demonstrate a set of mutually orthogonal binders operating simultaneously on one structure.

The macrophage experiments extend the approach from cargo attachment to receptor function. Across the 0-, 12- and 30-amino-acid *cis*-dimer constructs, all three receptor variants drove target-specific phagocytosis, with the 12- and 30-amino-acid linkers supporting robust engulfment alongside their IgG positive controls.

Accommodating a broad range of inter-domain linker strategies makes the receptor architecture forgiving in practice, offering latitude to incorporate diverse binders, extracellular geometries and signaling formats in future implementations. Sensitivity to ligand spacing on DNA origami pegboards also shows that DBP CARs can be paired directly with DNA-based spatial patterning to study phagocytosis. Because the DNA-binding domain is genetically encoded within the receptor polypeptide rather than added as an exogenous conjugate, newly expressed DBP CARs arrive at the cell surface inherently ligand-competent, supporting sustained phagocytic activity over a 24-hour interval after cell preparation under conditions where chemically labeled DNA CARs declined.

General-purpose addressing now depends mostly on the supply of binders. A six-base-pair site admits 4^6^, or 4,096, sequences; because a duplex and its reverse complement are the same site, 2,080 of these are distinct, which is ample space from which to draw mutually orthogonal pairs. Because specificities are generated computationally rather than found in nature, the set can be extended on demand, and the design principles established here provide a modular blueprint as new sequence specificities become available.

Binders need not remain restricted to canonical duplex sequences. The features that distinguish a nucleic acid nanostructure from genomic DNA — hairpin loops, crossover junctions, triplexes, DNA–RNA and DNA–PNA hybrid regions, locally strained duplexes, and chemically modified sites — are themselves recognizable, and a binder that reads geometry as well as sequence would reach positions that sequence-directed binding cannot. We expect surface-accessible docking of this kind to matter most where many different proteins must be placed on one structure, which is the regime in which conjugation chemistry becomes impractical and in which hybrid protein–DNA nanostructures have so far been hardest to build.

The common attachment strategy in the origami and bead experiments is recognition of a DNA sequence by a genetically encoded protein domain. DNA-functionalized materials therefore offer potential substrates beyond origami, provided that the recognition sites are accessible. Protein cargo can be expressed together with its binding domain, avoiding a separate protein–oligonucleotide conjugation step at the docking interface. Because the same D4 adapter accommodates fluorescent proteins, enzymes, antibody fragments and modular adaptors such as SpyCatcher, it can be extended to secondary recruitment — including nanoparticles, multi-enzyme cascades or additional catalytic readouts — without re-engineering the DNA-binding interface.

## Methods

### Materials and oligonucleotides

All oligonucleotides were purchased from Integrated DNA Technologies (IDT). Strands for hairpin, duplex and immobile crossover junction (ICJ) assays were ordered with HPLC or PAGE purification. Staple strands for Q1 DNA origami were ordered in 96-well V-bottom plates at standard desalting grade and resuspended to 200 μM in 1× TE buffer (pH 7.5). The p3364-DBP35opt scaffold strand (phagemid Addgene #266073) was produced using the eScaf *E. coli* strain (Addgene #220921) and purified in-house as previously described.^53^ All DNA sequences, Q1 origami scaffold and staple sequences, protein construct lists, buffer compositions and BLI kinetics are provided in the Supplementary Tables. Plasmids encoding the DNA-binding proteins (DBPs) were synthesized by GenScript in pET24a(+), pET30a(+) or pET22b(+) expression vectors and deposited with Addgene (IDs 262858– 262886).

Precast Novex TBE polyacrylamide gels (6% and 20%; Thermo Fisher Scientific) were run in XCell SureLock Mini-Cell systems using Ultra-Low Range (Fisher Scientific, 10-597-012), 50 bp (Fisher Scientific, 10-416-014) or 1 kb (NEB, N3232L) DNA ladders. DNA gels were stained with SYBR Safe (precast agarose, 8 μL per gel) or post-stained with SYBR Gold (2 μL in 20 mL buffer). SDS–PAGE (160 V, 35 min, room temperature) and native protein PAGE (100 V, 2 h, ice bath at 4 °C) used precast Any kD Mini-PROTEAN TGX gels (Bio-Rad) stained with InstantBlue Coomassie Protein Stain (Abcam, ISB1L).

### Protein expression and purification

Selected DBP constructs were expressed and purified by GenScript; in-house preparations followed a previously reported protocol^48^ with minor modifications. Plasmids were transformed into *Escherichia coli* BL21(DE3*) cells (NEB, C2527H) with a 30 s heat shock at 42 °C and plated on LB agar containing 50 μg mL^−1^ kanamycin (Teknova, L5025) overnight (>16 h) at 37 °C. Primary cultures (100 mL LB with 50 μg mL^−1^ kanamycin) were grown overnight (∼16 h) at 37 °C, and 20 mL was used to inoculate 700 mL LB with 50 μg mL^−1^ kanamycin.

Cultures were grown at 37 °C with shaking at 200 rpm to OD_600_ = 0.5–0.6, induced with 0.5 mM IPTG, and expressed for 4 h at 37 °C. Cells were harvested (3,500× *g*, 20 min, 4 °C), flash-frozen in liquid nitrogen and stored at −80 °C.

Pellets were thawed in lysis buffer (Supplementary Tables) and lysed on ice by sonication (35% amplitude, 5 s on / 10 s off for 10 min; three cycles with 10 min intervals). Lysates were clarified twice by centrifugation (29,000× *g* at 4 °C for 30 min, then 10 min), passed through a 0.33 μm syringe filter, and incubated with equilibrated Ni-affinity resin for 1 h or overnight at 4 °C. Resin was washed 4–5 times with 50 mL NiB1 buffer and eluted in 20 mL aliquots of NiB2 buffer. Pooled eluates (∼60 mL plus 10 mL IEB1) were loaded onto a HiTrap Heparin HP column (Cytiva, 17040701) on an ÄKTA pure system, washed with 7 column volumes (CV) of IEB1, and eluted with IEB2 (0–60% over 20 CV, 60–100% over 2 CV, hold at 100% for 2 CV). Relevant fractions (tracked by SDS –PAGE and, where applicable, 280 nm versus 488 nm for EGFP or 551 nm for mKO2) were pooled, concentrated using centrifugal MWCO filters at 4 °C, filtered through a 0.22 μm spin filter (Sigma-Aldrich, UFC30GV0S; 5,000× *g*, 2 min, 4 °C), and polished by size-exclusion chromatography on a Superdex 200 Increase 10/300 GL column in 1× FOB containing 14 mM MgCl_2_.

### DNA nanostructure assembly and purification

Hairpins and duplexes were assembled at 4 μM per strand in 1× FOB with 14 mM MgCl_2_ in a Bio-Rad DNA Engine Tetrad 2 thermocycler by holding at 90 °C for 5 min and cooling to 23 °C in 0.5 °C steps over approximately 1 h. Immobile crossover junctions (ICJs) were annealed identically at 4 μM per strand in 60 μL, resolved on 6% TBE gels (100 V, 150 min, ice bath at 4 °C), and excised from unstained sample lanes using flanking SYBR Safe-stained control lanes as guides. Excised gel slices were eluted overnight at 4 °C in 500 μL 1× FOB (14 mM MgCl_2_) in Freeze ‘N Squeeze tubes, centrifuged at 2,000× *g* (4 °C), concentrated with 30 kDa MWCO filters, and quantified by *A*_260_ (NanoDrop).

Q1 DNA origami was folded in 100 μL reactions containing 20 nM p3364-DBP35opt scaffold and 200 nM of each staple in 1× FOB with 4 mM MgCl_2_ (65 °C for 5 min; 60 °C to 40 °C at 0.5 °C per 30 min; cooled to 23 °C; one scaffold batch was folded at 12 mM MgCl_2_; Supplementary Fig. S7a). Folded origami was purified and buffer-exchanged into 1× FOB containing 14 mM MgCl_2_ by five washes on 100 kDa MWCO centrifugal filters (0.5 mL, pre-washed with 1× FOB + 14 mM MgCl_2_; 2,000× *g*, 20 min, 23 °C per wash) and recovered by inverted centrifugation for 5 min.

### Binding, gel-shift and enzyme-activity assays

Unless noted otherwise, proteins and DNA substrates were mixed at room temperature and incubated for at least 10 min before analysis. Hairpin assays used a 10-fold molar excess of DBP35opt or DBP48. For immobile crossover junction assays (Fig. 2b, c), junctions carrying a 5′-Cy5.5-labeled strand (sequences in Supplementary Tables) were incubated with mKO2–DBP35opt and resolved by native TBE PAGE; for display in Fig. 2c, the red fluorescence channel of both gel segments was uniformly gamma-adjusted (R_out_ = 255× (R_in_/255)^0.55^), while green and blue channels and Fig. 2b were unmodified. For *cis*-dimer binding to 34-bp duplexes, pre-annealed duplexes (8 μM, 2.5 μL) were mixed with a fourfold molar excess of protein and analyzed by 6% native TBE PAGE (0.5× TBE, 11 mM MgCl_2_, 100 V, 120 min, ice bath). For thermal annealing tests, mixtures were heated to 37, 40 or 55 °C for 10 min and cooled to 23 °C for 10 min prior to PAGE. Selected 34-5 duplex assays were also performed in 1× PBS with 14 mM MgCl_2_ and in cell binding buffer.

For Q1 origami binding, proteins were added at 6-, 12-, 24- or 48-fold molar excess relative to origami (reported as input protein molecules per origami scaffold). Q1 concentrations were 15 nM for agarose gel electrophoresis unless noted otherwise and 5 nM for negative-stain TEM; samples were incubated for 10 min at room temperature or overnight at 4 °C. The mScarlet3 and SpyCatcher gel assays in Fig. 5f, g used 20 nM Q1 in 15 μL and overnight incubation at 4 °C. For SpyCatcher–SpyTag recruitment, SpyCatcher3–D4 and SpyTag3–MBP– mScarlet3 were pre-reacted overnight at a 1:1.2 molar ratio before overnight incubation with Q1 at 4 °C. Origami Mg^2+^ screens used 2% agarose gels (0.5× TBE, 11 mM MgCl_2_, 90 V, 2 h). D4–cargo binding assays used 1.5% agarose gels (0.5× TBE, 11 mM MgCl_2_, ice bath at 4 °C; 60 V for 3 h for mScarlet3 and alkaline phosphatase, 5 h for SpyCatcher–SpyTag) post-stained with SYBR Gold for 30–60 min. For alkaline phosphatase (D4–ALP) assays, EDTA-free 0.2 M Tris-HCl, 10 mM MgCl_2_ and 1 mM ZnCl_2_ was used for gel casting and running (with D4–ALP diluted in the same buffer plus 200 mM NaCl); after SYBR Gold imaging, enzymatic activity was detected by incubating the gel for 60 min in 1-Step NBT/BCIP Substrate Solution (Thermo Scientific, 34042).

### Biolayer interferometry

Binding kinetics were measured on an Octet RED384 instrument (Sartorius) using streptavidin (SA) biosensors in PBS supplemented with 0.05% Tween-20 and 0.1% BSA. Biotinylated DNA constructs (20 nM) were immobilized to a loading response of ∼0.35 nm and dipped into a five-point, threefold analyte dilution series (500, 167, 56, 19 and 6 nM) alongside a buffer reference well, with 5 min association and 5 min dissociation phases.

The kinetic parameters reported for all six assays in Supplementary Table S1 were obtained from global 2:1 (heterogeneous ligand) fits in Octet Analysis. For the 30AAFL target-DNA series, this analysis yielded an apparent K_D_ of 0.124 nM and k_off_ of 4.25× 10^−5^ s^−1^ for the slow-dissociating component. Processed sensorgrams, fit exports and analysis scripts are provided in the BLI data package.

### Computational design of *trans*-dimers

Target recognition-site coordinates on helices 8 and 10 (scaffold residues 1346–1351 and 1417–1422) were taken from an atomic model generated from the Q1 caDNAno design using tacoxDNA,^54^ preserving their native spacing and relative orientation in the folded array. Using PyMOL (Schrödinger, LLC), two copies of the 63-residue DBP35opt monomer were positioned onto the two target DNA sites (one copy at each site), and the DNA chains and remainder of the origami were omitted for subsequent protein-only backbone generation. Two-site constructs were built by circular permutation, opening the first DBP35opt chain between residues 39 and 40 to insert the second complete copy flanked by *de novo* connecting segments (20–100 residues) and terminal extensions (10–50 residues). In D4 (327 residues), these regions comprise 22, 49, 90 and 39 residues, respectively (127 parent-derived and 200 designed residues). Holding the two positioned DBP35opt protein modules fixed, backbones (∼1,000 per condition) were generated with RFdiffusion (v1)^55^ to build the connecting segments and terminal extensions fusing both modules into a single polypeptide. Sequences for generated regions were designed with ProteinMPNN^56^ while holding parent DNA-contacting residues fixed (without nucleic acid context). Fold self-consistency was evaluated by predicting standalone protein structures with AlphaFold 3^57^ (default parameters) and scoring predicted confidence and full-construct backbone RMSD against the design model. Four designs (D4, D10, D34 and D46) were selected for expression; D4 achieved pTM = 0.93, mean pLDDT = 88.3 and backbone RMSD < 0.9 Å.

### Negative-stain TEM and cryo-EM

For negative-stain TEM, Formvar/carbon-coated 400-mesh copper grids (Ted Pella, 01754-F) were glow-discharged (PELCO; 30 s hold, 30 s discharge at 15 mA, 5 min rest). Grids were floated on 3 μL of Q1 sample for 2 min, washed on two 8 μL drops of water (brief, then 10 s), stained on two 3 μL drops of 0.75% uranyl formate (brief, then 1 min), air-dried, and imaged on a Tecnai T12 microscope (FEI) at 30,000× or 45,000× nominal magnification.

For cryo-EM of the ds34-2 / no-linker *cis*-dimer complex (3 μM ds34-2 DNA, 3 μM H8TEV–EGFP– 2×DBP35opt), vitrified grids were imaged on a Glacios transmission electron microscope (Thermo Fisher Scientific; 200 kV, *C*_s_ = 2.7 mm, Gatan K3 detector, 54,000× nominal magnification, 0.73 Å pixel^−1^, total dose 60.0 e^−^ Å^−2^). Across 1,177 imported motion-corrected micrographs processed in cryoSPARC^58^ (v4.3.1) with Patch CTF estimation, 646 exposures passed curation. Blob picking (20–116 Å diameter) identified 2,477,188 particles, of which 2,102,590 were extracted in a 400-pixel box Fourier-cropped to 100 pixels (2.92 Å pixel^−1^). Iterative 2D classification and selection retained 99,240 particles, and 98,401 were re-extracted in the full 400-pixel box (0.73 Å pixel^−1^). Following three-class *ab initio* reconstruction, duplicate removal (85,226 particles) and three rounds of three-volume heterogeneous refinement (retaining 28,224, 15,436 and 9,505 particles), 2D classification aligned 8,512 particles and 2D class selection retained 8,190 particles for the final three-class *ab initio* reconstruction (2,899, 2,050 and 3,241 particles per class) and non-uniform refinement.

For cryo-EM of the ds34-5 / 30AAFL *cis*-dimer complex (3 μM ds34-5 DNA, 1.5 μM H8TEV–EGFP– 2×DBP35opt–30AAFL; heated at 55 °C for 10 min and cooled to 23 °C for 10 min), 3.2 μL was applied to glow-discharged Quantifoil R1.2/1.3 300-mesh gold grids and plunge-frozen on a Vitrobot (Thermo Fisher Scientific; 23 °C, 90% humidity, blot force 0, 2–4 s blot). A total of 14,724 movies were collected on a Titan Krios (Thermo Fisher Scientific; 300 kV, *C*_s_ = 2.7 mm, Gatan K3 detector, 105,000× nominal magnification, 0.8189 Å pixel^−1^, total dose 47.7 e^−^ Å^−2^) and preprocessed in cryoSPARC Live (v5.0.2) with Patch Motion Correction and Patch CTF estimation. In cryoSPARC (v5.0.4), template picking (140 Å diameter) using 50 projections of a theoretical model volume yielded 20,406,737 extracted particles (288-pixel box Fourier-cropped to 72 pixels, 3.2756 Å pixel^−1^). Four rounds of 200-class 2D classification and selection retained 5,136,003 particles upon re-extraction in a 280-pixel box (0.8189 Å pixel^−1^). Seven-class *ab initio* reconstruction, seven-volume heterogeneous refinement (2,131,621 particles), five-class *ab initio* reconstruction (1,024,298 particles) and two-class *ab initio* reconstruction isolated a 509,960-particle population; subsequent 2D classification retained 509,016 aligned particles for the final three-class *ab initio* reconstruction (140,343, 146,066 and 222,607 particles per class).

For the monomeric Q1 control (Q1 origami incubated with a fivefold molar excess of H8TEV–EGFP– DBP35opt), 5,384 motion-corrected micrographs were collected on a Glacios (200 kV, *C*_s_ = 2.7 mm, K3 detector, 0.73 Å pixel^−1^, total dose 60.0 e^−^ Å^−2^) and processed in cryoSPARC (v4.3.1) with Patch CTF estimation, retaining 5,076 exposures. Initial blob picking (300–560 Å) and 2D classification generated templates for template picking (560 Å diameter, 287,533 particles extracted at 1,024 pixels Fourier-cropped to 64 pixels), followed by 2D curation, full-box re-extraction (1,024 pixels, 10,552 particles) and 2D classification into three final classes (5,231 particles).

For Q1 with D4 (15 nM origami with 180 nM D4, 12× molar excess, 10 min at room temperature), 3.2 μL was applied to in-house amine-functionalized graphene oxide grids^59^ and vitrified on a Vitrobot (23 °C, 90% humidity, blot force −1, 2–6 s blot). Movies were collected on a Titan Krios (300 kV, *C*_s_ = 2.7 mm, Gatan K3 detector, 105,000× nominal magnification, 0.8189 Å pixel^−1^, total dose 47.7 e^−^ Å^−2^) and processed in cryoSPARC Live (v5.0.2) with Patch Motion Correction and Patch CTF estimation across 11,265 picked exposures (11,263 exported accepted exposures). Circular blob picking (100–560 Å diameter) yielded 6,693,044 exported particles extracted in a 1,024-pixel box Fourier-cropped to 64 pixels (13.10 Å pixel^−1^). Following streaming and offline 2D classification, selection and duplicate removal in cryoSPARC (v5.0.4), 121,055 particles were re-extracted in the full 1,024-pixel box (0.8189 Å pixel^−1^). One round of 2D classification and selection (70,266 particles), duplicate removal at 50 Å minimum separation (59,273 particles), and two rounds of 2D classification and selection (57,717 aligned particles reduced to 34,368 and 33,974 particles) yielded 33,768 aligned particles across 50 classes in the final 2D classification (the class average shown in Fig. 5d contains 6,042 particles).

### Cell culture, receptor constructs and phagocytosis assays

RAW264.7 macrophages (ATCC TIB-71; <20 passages) were cultured in DMEM with 10% FBS, 1% penicillin– streptomycin–glutamine (PSG) and 1 mM sodium pyruvate at 37 °C and 5% CO_2_. HEK293T cells (Takara, 632180) were cultured in DMEM with 10% FBS and 1% PSG for lentivirus production (pMD2.G, Addgene #12259; pCMV-dR8.2, Addgene #8455; Lipofectamine LTX, Invitrogen 15338-100; concentrated with Lenti-X, Takara 631232).

DBP CAR variants were cloned into pHR lentiviral vectors encoding, from N to C terminus: CD3ε signal peptide, HA tag, LPETGGGGGG linker, tandem DBP35opt *cis*-dimer (0-, 12- or 30-aa flexible linker), GGSGGSGGS linker, CD86 transmembrane domain, mouse FcεRIγ cytoplasmic domain (aa 45–86, UniProt P20491), GSGS linker and EGFP. The activating FcγR DNA CAR control (Addgene #176611) carried SNAPf in place of the DBP dimer and mGFP5 at the C terminus.

Small unilamellar vesicles (SUVs; 97.8% POPC, 2% biotinyl cap PE, 0.1% PEG5000-PE, 0.1% Atto647-DOPE) were prepared by freeze–thaw cycles, diluted to 2 mM in PBS (pH 7.2) and 0.22 μm filtered. Silica beads (5.01 μm; Bangs Labs, SS05003) were coated with 1 mM SUVs (30 min), blocked with 0.2% casein in PBS (15 min), and incubated with 1 μg mL^−1^ neutravidin (20–30 min). Beads were functionalized for 1 h in PBS with 20 mM MgCl_2_ using 250 nM biotinylated target or control duplex DNA, 250 nM biotinylated ssDNA ligand (DNA CAR control), or saturating amounts of PEG-purified DNA origami pegboards (p8064 scaffold, 8T tight or 8S sparse ligand patterns, folded and purified as described^49^; sequences in Supplementary Tables).

For phagocytosis assays, ∼8× 10^4^ RAW264.7 macrophages were plated per well in 96-well glass-bottom plates (Brooks, MGB096-1-2-LG-L) ∼24 h prior to assay. For DNA CAR controls, benzylguanine-conjugated receptor DNA (prepared via overnight reaction of 5′-amine oligo with BG-GLA-NHS at 1:50 molar ratio and NAP-5 purification^49,60^) was incubated with cells at 1 μM in serum-free DMEM for 10 min and washed out; DBP CAR cells required no labeling. Approximately 8× 10^5^ functionalized beads were added per well for 30 min at 37 °C. Cells were imaged by spinning-disc confocal microscopy (40×, 0.95 NA air objective), and internalized beads were scored in ImageJ by a blinded analyzer (Blind-Analysis-Tools-1.0), with ≥100 cells scored per independent experiment.

### Statistical analysis

Phagocytosis data are summarized as means ± SEM, with n = 4 experiments per condition in Fig. 4b, c, e and n = 3 per condition in Fig. 4d. At least 100 cells were scored per independent experiment. Figure 4b used repeated-measures one-way ANOVA with Geisser–Greenhouse correction and Tukey-adjusted comparisons across the four conditions. Figure 4c used repeated-measures two-way ANOVA, matching both ligand condition and linker variant, with Geisser–Greenhouse correction and Tukey adjustment across all 66 pairwise comparisons among the 12 condition means. Figure 4d used ordinary two-way ANOVA with Šídák adjustment across all 28 pairwise comparisons among the eight condition means. Figure 4e was normalized within each experiment and condition to its 1-hour value; conditions were matched by experimental day. Reported adjusted P values are from the saved Prism analyses.

Internal-target-zone occupancy in the four-design negative-stain TEM screen was quantified by manual particle counting across micrographs (24-fold molar excess for D4 and D34; sixfold for D10 and D46). Original micrographs and manual annotations are provided in Supplementary Data 1.

### Structural models and visualization

The *trans*-dimer design workflow used PyMOL for two-site DBP35opt placement on the Q1 target helices, RFdiffusion (v1)^55^ for protein-only backbone fusion and ProteinMPNN^56^ for sequence design. AlphaFold 3^57^ predictions of the standalone protein sequences were used to assess fold self-consistency and render the 327-residue D4 structure shown in Fig. 5e and Supplementary Fig. S8.

The DBP35opt structure in Supplementary Fig. S8 was extracted from the EGFP–DBP35opt coordinate model. Parent and D4 coordinates were rendered in UCSF ChimeraX 1.12.^61^ The parent was aligned to D4 domain 1 by a least-squares fit of 60 corresponding Cα positions. The second view of each protein was generated by a 90° rotation about the vertical axis. Colors identify corresponding parent-residue positions; gray denotes *de novo* designed D4 bridge and extension residues.

Figure 5e shows the AlphaFold 3 D4 prediction modeled onto 18-bp recognition-site duplex segments of helices 8 and 10 from the designed Q1 atomic array by least-squares superposition of the conserved DBP35opt–DNA recognition motif onto both D4 domains followed by restrained all-atom clash relief with fixed DNA coordinates, rendered in UCSF ChimeraX 1.12.^61^ The Q1 representations in Fig. 5a–c were derived from the same designed Q1 atomic model.

### Use of large language models

During manuscript preparation, the authors used Codex GPT-6-Astra and Gemini 4 Neon to assist with language editing, formatting, and scripts for data organization and figure rendering. The authors reviewed and verified all resulting text and code and take full responsibility for the content of the manuscript.

## Acknowledgments

We thank Wooyoung Choi, Nitesh Khandelwal, Dominic Grisingher, Thalappil Pradeep, and Sanchari Saha for helpful discussions. We thank Zanlin Yu, Kevin Choi, and Adamo Mancino for support in making amino-functionalized graphene-oxide grids and protocols for early tests. We thank David Bulkley and Glenn Gilbert for help with cryo-EM data collection and interpretation.

## Funding

This work was supported by the National Institutes of Health under awards R35GM125027 (to S.M.D.) and R35GM146935 (to M.M.). M.M. is a Nadia’s Gift Foundation Innovator of the Damon Runyon Cancer Research Foundation (DRR-85-25). Cryo-EM equipment at the UCSF EM Core is partially supported by NIH grant S10OD026881.

## Author contributions

Conceptualization: R.P.N., S.M.D. Methodology: R.P.N., E.S., A.P., C.J.G., K.S. Software: A.P., C.J.G. Investigation: R.P.N., E.S., O.G., S.R., K.S. Formal analysis: R.P.N., E.S., M.M., S.M.D. Resources: K.S., C.J.G. Visualization: R.P.N., E.S. Supervision: S.M.D., M.M., C.J.G. Funding acquisition: S.M.D., M.M., C.J.G. Project administration: R.P.N., S.M.D. Writing – original draft: R.P.N., E.S. Writing – review and editing: R.P.N., O.G., E.S., A.P., C.J.G., M.M., S.M.D.

## Competing interests

The authors declare no competing interests.

## Data availability

Plasmids encoding the DNA-binding proteins are available from Addgene under accession numbers 262858–262886, and the custom p3364-DBP35opt scaffold phagemid (produced using the eScaf strain, Addgene #220921) under accession number 266073. DNA and protein sequences, Q1 origami staple sequences, buffer compositions and BLI kinetics are provided in the Supplementary Tables. DNA nanostructure design files, supplementary gels and related images, negative-stain TEM micrographs, supporting assay and construct data, and computational models are deposited in Zenodo (https://doi.org/10.5281/zenodo.24519036).^62^ Cryo-EM 2D class averages and *ab initio* volumes are available from the corresponding author upon reasonable request.

## Code availability

Custom analysis and figure-rendering scripts, together with generic AlphaFold 3 input-preparation and inference utilities, are available in Zenodo (https://doi.org/10.5281/zenodo.24519036)^62^ under the MIT License (with the bundled AlphaFold 3 runner retaining its CC BY-NC-SA 4.0 license and excluding model weights). DNA nanostructure design files are also hosted at https://github.com/douglaslab/cadnano-designs. The original project-specific RFdiffusion and ProteinMPNN pipeline scripts were not retained; the D4 design model and PyMOL session are archived with the supporting data.

## Notes

https://doi.org/10.5281/zenodo.24519036

### Competing Interest Statement

The authors have declared no competing interest.

### Summary of Updates

The updated version corrects author names, funding sources, formatting, and adds an Addgene accession number and AI use disclosure.

