## Supplementary Information for "Protein docking on DNA surfaces using *de novo* designed binders"

S1. Materials and oligonucleotides
S2. Additional purification and DNA assembly details
S3. Protein–DNA binding and computational design
S4. Additional electron microscopy details
S5. Biolayer interferometry
S6. Additional phagocytosis assay details
S7. Supplementary figures, table and sequence files
S8. References

### S1. Materials and oligonucleotides

Materials, oligonucleotide sources and gel electrophoresis conditions are described in Methods, "Materials and oligonucleotides." Scaffold production (p3364-DBP35opt phagemid, Addgene #266073) followed the published eScaf protocol.[1] DNA sequences and protein construct lists are provided in Supplementary_Tables.xlsx; buffer compositions are retained in Tables 1 and 2 below. Oligonucleotide preparation specific to the cell assays is described in Section S6.

### S2. Additional purification and DNA assembly details

Protein expression and purification followed the protocol described in Methods, "Protein expression and purification," adapted from Glasscock et al.[2] Additional handling details are given below.

Following transformation and recovery, 500 µL of culture was plated. A large colony mass was collected with a sterile swab to inoculate the primary culture. Harvested cell pellets were transferred to 50 mL conical tubes and centrifuged again before flash-freezing.

The lysis and purification buffer compositions are listed in Table 1.

**Table 1.** Protein purification buffers and their compositions.

| Buffer | HEPES | NaCl | Imidazole | TCEP | pH |
|---|---|---|---|---|---|
| **NiB1** | 25 mM | 150 mM | 5 mM | 0.2 mM | 8.0 |
| **NiB2** | 25 mM | 150 mM | 500 mM | 0.2 mM | 8.0 |
| **IEB1** | 25 mM | 150 mM | — | 0.2 mM | 8.0 |
| **IEB2** | 25 mM | 1 M | — | 0.2 mM | 8.0 |

**Lysis Buffer (100 mL):** 100 mL NiB1 buffer + 1 SigmaFAST protease-inhibitor tablet + pinch of lysozyme + DNase I (added at the end).

Lysates were clarified using a Sorvall Lynx 4000 centrifuge. Ni-affinity binding was performed with mixing; no appreciable difference in protein yield or subsequent purification was observed between 1 h and overnight incubations. During each wash, the resin was resuspended and allowed to settle for approximately 5 min before removing the wash buffer. Flow-through, wash and elution fractions were checked by SDS–PAGE.

Concentrated protein was spin-filtered using an Eppendorf 5425 F centrifuge and immediately loaded for size-exclusion chromatography. Separation proceeded over approximately 1.2–1.4 column volumes. SEC fractions were checked by SDS–PAGE before pooling for binding and biochemical assays.

**Table 2.** SEC and binding buffers and their compositions.

| Buffer | Tris-HCl | HEPES | $MgCl_2$ | NaCl | EDTA | $Na_2HPO_4$ | KCl | $KH_2PO_4$ | $CaCl_2$ | Glucose | pH |
|---|---|---|---|---|---|---|---|---|---|---|---|
| **FOB14** | 5 mM | — | 14 mM | 5 mM | 1 mM | — | — | — | — | — | 7.5 |
| **PBS14** | — | — | 14 mM | 137 mM | — | 10 mM | 2.7 mM | 1.8 mM | — | — | 7.5 |
| **CBB-20** | — | 20 mM | 20 mM | 135 mM | — | — | 4 mM | — | 1 mM | 10 mM | 7.5 |

Hairpin, duplex, ICJ and Q1 assembly, annealing and purification are described in Methods, “DNA nanostructure assembly and purification.” For ICJ recovery, excised gel pieces were fragmented and shaken overnight in elution buffer. After concentration, the 30 kDa MWCO filter was inverted and centrifuged for 2 min to recover the sample.

For the 15 µL, 20 nM Q1 gel samples, 3 µL of 6× loading dye was added before electrophoresis. Mixed complexes and unmixed controls containing excess protein were maintained at 4 °C before loading. Gel compositions, running conditions and ALP activity detection are described in Methods, “Binding, gel-shift and enzyme-activity assays.”

### S3. Protein–DNA binding and computational design

Protein–DNA incubation conditions, molar ratios, thermal annealing tests and buffer comparisons are described in Methods, “Binding, gel-shift and enzyme-activity assays.” Protein excess for Q1 assays is expressed per origami scaffold. Q1 contains 12 recognition sites, so a 12-fold input excess corresponds to one protein molecule per recognition site; this input ratio does not specify binding occupancy.

The design workflow is described in Methods, “Computational design of trans-dimers.” DBP35opt modules were positioned in PyMOL using the native Q1 target-site geometry; DNA was then omitted for protein-only backbone generation with RFdiffusion (v1). ProteinMPNN retained the parent DNA-contacting residues, and standalone AlphaFold 3 predictions assessed fold self-consistency.

The original project-specific RFdiffusion pipeline scripts were not retained after cluster storage cleanup. The retained D4 model and PyMOL session are provided with the supporting data; the available AlphaFold 3 scripts are generic utilities, not the original project-specific pipeline. The tested trans-dimers were D4, D10, D34 and D46 (deposited with Addgene as H8_TEV_DimerCameronD4_pET24a, H8_TEV_DimerCameronD10_pET24a, H8_TEV_DimerCameronD34_pET24a and H8_TEV_DimerCameronD48_pET24a, respectively; D46 carries the legacy label D48 in the Addgene deposit). Protein construct lists, sequences and Addgene plasmid names are provided in Supplementary_Tables.xlsx (Section S7).

### S4. Additional electron microscopy details

Grid preparation, microscope settings and dataset-specific cryo-EM processing are described in Methods, “Negative-stain TEM and cryo-EM.” The ds34-5–30AAFL and Q1–D4 datasets have separate particle-picking and classification workflows, as detailed there.

For negative staining, sample drops were placed on paraffin film and grids floated carbon-side down. Excess liquid was wicked away with Whatman filter paper after sample application and each wash and stain step. Grids were air-dried for at least 5 min and up to overnight before imaging.

For cryo-EM, both ds34-5–30AAFL and Q1–D4 samples were applied to glow-discharged grids with no wait time before blotting. Q1–D4 used in-house amine-functionalized graphene oxide grids prepared as described previously.[3] The distinct grid types and blot settings for the two samples are given in the main Methods.

### S5. Biolayer interferometry

Experimental conditions are described in Methods, “Biolayer interferometry.” The ligand panel included hairpin DNA, double-stranded DNA and control DNA. Supplementary Table S1 reports global 2:1 (heterogeneous ligand) fit parameters for all six assays. For the 30AAFL target-DNA series, the reported 0.124 nM value is an apparent affinity of the slow-dissociating component, whose precise value is poorly constrained. The saved vendor settings specify a one-component model, whereas the reported fit exports contain two components; the exact final vendor analysis state is unavailable. The original FRD file for sensor D3 (19 nM) is missing, but processed and tabular raw-data exports are retained. Sensorgrams, fit exports and sensitivity analyses are provided in the BLI data package.

### S6. Additional phagocytosis assay details

Cell culture conditions, receptor domain architecture, bead functionalization and assay timing are described in Methods, “Cell culture, receptor constructs and phagocytosis assays.” Additional sequences and preparation details are retained below.

#### Expressed Constructs

DBP CAR sequence details. The CD3ε signal peptide is MQSGTHWRVLGLCLLSVGVWGQD and the HA tag is YPYDVPDYA. The three variants differ in the inter-domain linker connecting the two DBP35opt copies: a 30-amino-acid flexible linker (GGSGGSGGSGGSGGSGGSGGSGGSGGSGGS), a 12-amino-acid flexible linker (GGSGGSGGSGGS), or no linker (0 aa, direct fusion). The DBP35opt domain sequence is: GFGRAVKEKRKELGLTQVEFAEKAGLSRRTIINIERGYIVPQKATKEKIAKALGTSVEELEQA. Construct coding sequences were codon-optimized for expression in mouse cells.

#### Oligonucleotides

**DBP CAR duplex ligand (5-base spacer).** The duplex ligand for the DBP CARs consists of two annealed oligonucleotides containing two TGCACA binding sites for DBP35opt separated by a 5-base spacer:

**35.dplxDimer.5f:** 5′-GC ATATAT TGCACA TTTTT TGCACA ATATATA GC/3Bio/-3′

**35.dplxDimer.5r:** 5′-GC TATATAT TGTGCA AAAAA TGTGCA ATATAT GC-3′

**Control duplex (no binding site).** A control duplex in which the TGCACA binding motifs were replaced with poly-T/poly-A runs, matched in length:

**35.Dimer.Cntrl_f:** 5′-GC ATATAT TTTTTT TATAT TTTTTT ATATATA GC/3Bio/-3′

**35.Dimer.Cntrl_r:** 5′-GC TATATAT AAAAAA ATATA AAAAAA ATATAT GC-3′

**DNA CAR receptor-ligand pair (control).** The activating FcγR DNA CAR receptor strand (5′-BG-/5AmMC6/AATATGATGTATGTGG-3′) was ordered from IDT with a 5′ terminal amine and conjugated to benzylguanine as described [4],[5]. The corresponding ligand strand (5′-/5BiosG/TTTT-TTTCATACATCATATT-3′) was ordered from IDT with a 5′ biotin.

All duplex oligonucleotides were purchased from IDT with standard desalting purification. Duplex ligands were annealed by mixing equimolar forward and reverse strands, heating to 95°C for 5 min, and cooling to room temperature.

#### DNA Origami Preparation

DNA origami pegboards were prepared as before[4]. Briefly, the p8064 DNA scaffold (IDT, Cat# 1081314) was mixed with core staple oligonucleotides (200 nM; plates 1 and 2), biotinylated oligonucleotides (200 nM), and fluorophore-labeled oligonucleotides (200 nM) in 1× folding buffer (5 mM Tris pH 8.0, 1 mM EDTA, 5 mM NaCl, 20 mM MgCl2) at a final scaffold concentration of 20 nM. For pegboards presenting DBP CAR ligands, handle staple oligonucleotides at ligand positions were extended at their 3′ end with the 35.dplxDimer.5f sequence plus a 5T spacer (200 nM each). The complementary reverse strand (35.dplxDimer.5r) was co-folded in the annealing reaction at 10× excess relative to each handle position (e.g. 1.6 µM total for 8-handle designs). Blank (no-ligand) positions used handle oligonucleotides from the Plate 3-L (no ligand) set. Folding was performed in a PCR thermocycler with initial denaturation at 65°C for 15 min followed by cooling from 60°C to 40°C at 1°C/hr. Excess oligonucleotides were removed by two rounds of PEG precipitation: the folding reaction was mixed with an equal volume of PEG precipitation buffer (15% w/v PEG-8000, 5 mM Tris pH 8.0, 1 mM EDTA, 500 mM NaCl, 20 mM MgCl2) and centrifuged at 16,000× rcf for 25 min at room temperature. The pellet was resuspended in 1× folding buffer after each round. Final pellets were resuspended at the desired concentration and stored at 4°C. Two origami designs were used: 8 tightly spaced ligands (8T) and 8 distantly spaced ligands (8S), as defined in Kern et al.[4] All staple and ligand sequences are listed in Supplementary_Tables.xlsx.

#### Supported Lipid Bilayer-Coated Beads

SUV preparation. Lipid identities and proportions are given in the main Methods: POPC (Avanti, 850457), biotinyl cap PE (Avanti, 870273), PEG5000-PE (Avanti, 880230) and Atto647-DOPE (ATTO-TEC, AD 647-161). Chloroform-suspended lipids were mixed and desiccated overnight. Lipid sheets were resuspended at 10 mM in PBS, pH 7.2 (Gibco, 20012050), and stored under nitrogen. After freeze–thaw cycles, SUVs were stored at −80 °C under nitrogen. Immediately before use, diluted SUVs were filtered through a 0.22 µm filter (Millipore, SLLG013SL).

Bead preparation. Silica beads were supplied at 10% solids and washed with PBS before coating. SUV coating and DNA functionalization were performed at room temperature with end-over-end mixing, with PBS washes between coating, casein blocking (Sigma, C5890), neutravidin incubation (Thermo, 31000) and ligand addition. Duplex-functionalized beads were washed and resuspended in PBS containing 0.2% casein and 20 mM $MgCl_2$. Origami pegboards were also added in PBS containing 0.2% casein and 20 mM $MgCl_2$, with end-over-end mixing during the 1 h incubation.[4]

#### Lentivirus Production and Infection

Lentivirus was produced using the plasmids and reagents listed in the main Methods. pMD2.G and pCMV-dR8.2 were gifts from Didier Trono and Bob Weinberg, respectively. Medium was harvested 72 h after transfection, passed through a 0.45 µm filter (Millipore, SLHVM33RS), and concentrated with Lenti-X. Concentrated virus was added directly to RAW264.7 macrophages.

#### Phagocytosis Assay

Internalized beads were identified by their fluorescent supported lipid bilayer. Imaging used a 40×, 0.95 NA Plan Apo air objective; blinded scoring and the number of cells scored are described in the main Methods.

#### Preparation of Benzylguanine-Conjugated DNA

For DNA CAR control experiments, benzylguanine-conjugated receptor DNA was prepared as previously described[4,5]. Briefly, 5′-amine-modified (5AmMC6) receptor oligonucleotides (IDT) were diluted to 2 mM in 0.15 M HEPES pH 8.5 and reacted overnight at room temperature with BG-GLA-NHS (NEB, Cat# S9151S) at a 1:50 amine:NHS molar ratio. Excess benzylguanine was removed using NAP-5 columns (Cytiva, Cat# 17085301), and the product was concentrated, resuspended in water at 100 µM, aliquoted, and stored at −20°C.

### S7. Supplementary figures, table and sequence files

#### Supplementary Table S1

BLI binding kinetics for monomeric and cis-dimeric constructs. NSB = Non-specific Binding

| Assay | Description | $K_d$ | $K_{on}$ | $K_{off}$ | NSB $K_d$ | NSB $K_{on}$ | NSB $K_{off}$ | Total R2 of curve |
|---|---|---|---|---|---|---|---|---|
| **Assay 1** | Single-stranded DNA hairpin with monomer protein | 1.74E-08 | 1.22E+05 | 2.12E-03 | 3.33E-07 | 8.95E+04 | 2.98E-02 | 0.997 |
| **Assay 2** | Single stranded DNA hairpin with mKO2-monomer fusion protein | 1.97E-08 | 3.18E+04 | 6.27E-04 | 2.43E-05 | 3.03E+03 | 7.36E-02 | 0.9953 |
| **Assay 3** | Double-stranded DNA with 30AA dimer protein | 1.24E-10 | 3.42E+05 | 4.25E-05 | 5.21E-07 | 9.08E+04 | 4.73E-02 | 0.9756 |
| **Assay 4** | Double-stranded DNA with 12 AA dimer protein | 1.50E-09 | 2.03E+05 | 3.04E-04 | 2.13E-06 | 1.29E+04 | 2.75E-02 | 0.987 |
| **Assay 5** | Control DNA with 30 AA dimer protein | 3.71E-09 | 2.74E+05 | 1.02E-03 | 4.44E-07 | 9.23E+04 | 4.10E-02 | 0.9923 |
| **Assay 6** | Control DNA with 12 AA dimer protein | 1.72E-09 | 1.79E+05 | 3.08E-04 | 1.66E-07 | 1.22E+05 | 2.02E-02 | 0.9922 |

#### Supplementary sequence files

All DNA sequences, Q1 origami scaffold and staple sequences, protein construct lists, buffer compositions and BLI kinetics are provided in Supplementary_Tables.xlsx. DNA nanostructure design files (caDNAno) and custom scaffold sequences are available at https://github.com/douglaslab/cadnano-designs and in Zenodo (https://doi.org/10.5281/zenodo.24519036) together with supporting images, data and code.

**a**

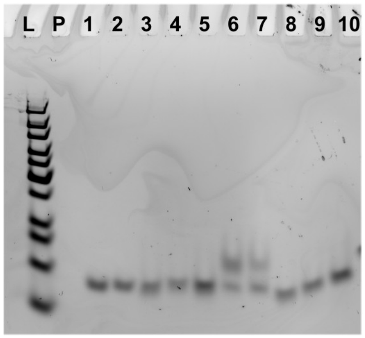


**L**-Ultra Low range Ladder
**P**- H8TEV_DBP35opt abbreviated as P35

**1**-T1_null.hp.1
**2**-T1_null.hp.2
**3**-P35 + T1_null.hp.1
**4**-P35 + T1_null.hp.2

**5**-P35+ T1_35.hp.1.
**6**-P35+ T1_35.hp.2.
**7**-P35 + T1_35.hp.3.

**8**-P35 + T1_48.hp.1.
**9**-P35 + T1_48.hp.2.
**10**-P35 + T1_48.hp.3.

**b**

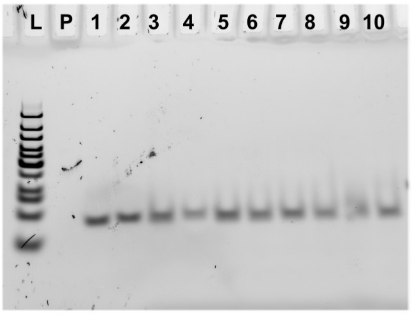


**L**-Ultra Low range Ladder
**P**- H8TEV_DBP48 abbreviated as P48

**1**-T1_null.hp.1
**2**-T1_null.hp.2
**3**-P48 + T1_null.hp.1
**4**-P48 + T1_null.hp.2

**5**-P48+ T1_35.hp.1.
**6**-P48+ T1_35.hp.2.
**7**-P48 + T1_35.hp.3.

**8**-P48 + T1_48.hp.1.
**9**-P48+ T1_48.hp.2.
**10**-P48+ T1_48.hp.3.

**c**

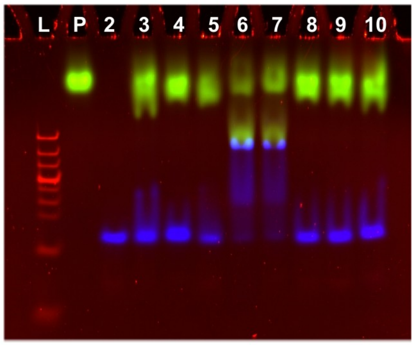


**L**-Ultra Low range Ladder
**P**- H8TEV_EGFP_DBP35opt abbreviated as E_DBP35

**2**-T1_null.hp.2
**3**-E_DBP35 + T1_null.hp.1
**4**-E_DBP35 + T1_null.hp.2

**5**-E_DBP35+ T1_35.hp.1.
**6**-E_DBP35+ T1_35.hp.2.
**7**-E_DBP35 + T1_35.hp.3.

**8**-E_DBP35 + T1_48.hp.1.
**9**-E_DBP35+ T1_48.hp.2.
**10**-E_DBP35+ T1_48.hp.3.

**d**

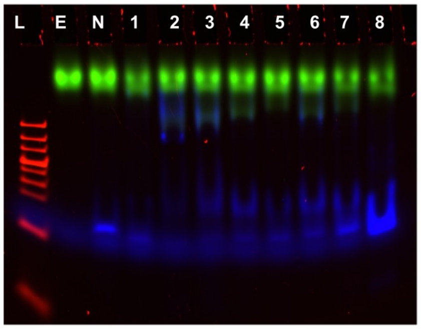


**L**-Ultra Low range Ladder
**E**- H8TEV_EGFP_DBP35opt abbreviated as E_DBP35

**C**-E_DBP35+ Null-hp4

**1**-E_DBP35 + 35dplx_1
**2**-E_DBP35 + 35dplx_2
**3**-E_DBP35 + 35dplx_3
**4**-E_DBP35 + 35dplx_4
**5**-E_DBP35 + 35dplx_5
**6**-E_DBP35 + 35dplx_6
**7**-E_DBP35 + 35dplx_7
**8**-E_DBP35 + 35dplx_8

**e**

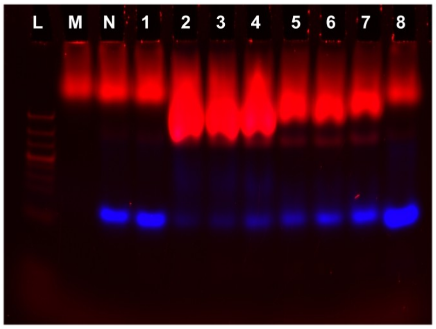


**L**-Ultra Low range Ladder
**M**-H8TEV_mKO2_DBP35opt abbreviated as M_DBP35

**C**-M_DBP35+ Null-hp4

**1**-M_DBP35 + 35dplx_1
**2**-M_DBP35 + 35dplx_2
**3**-M_DBP35 + 35dplx_3
**4**-M_DBP35 + 35dplx_4
**5**-M_DBP35 + 35dplx_5
**6**-M_DBP35 + 35dplx_6
**7**-M_DBP35 + 35dplx_7
**8**-M_DBP35 + 35dplx_8

**f**

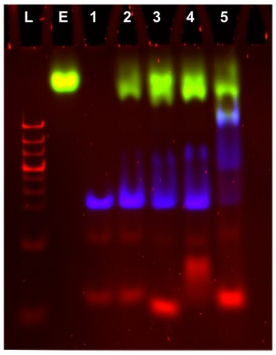


**L**-Ultra Low range Ladder
**E**- H8TEV_EGFP_DBP35opt abbreviated as E_DBP35
**C**-Null-dplx_1
**1**-E_DBP35opt+ Nulldplx_1
**2**-E_DBP35 + Nulldplx_2
**3**-E_DBP35 + 48-dplx_1
**4**-E_DBP35 + 35dplx_1

**Supplementary Figure S1. Monomer and fluorescent fusion binding to hairpin and duplex DNA.** (a–c) Hairpin binding by DBP35opt (a), DBP48 (b) and EGFP–DBP35opt (c). (d, e) Cy5.5-labeled hairpins with EGFP–DBP35opt (d) and mKO2–DBP35opt (e). (f) Cy5.5-labeled duplexes with EGFP–DBP35opt. Lane identities are listed beneath each gel. DNA in c–f carries a 5′ Cy5.5 label. Sequences are provided in Supplementary_Tables.xlsx.

**a** Before heating

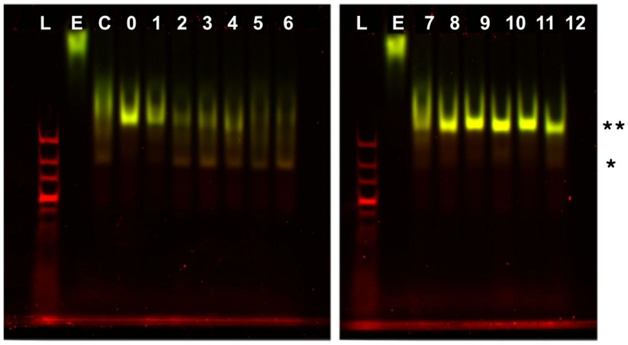


**b** After heating

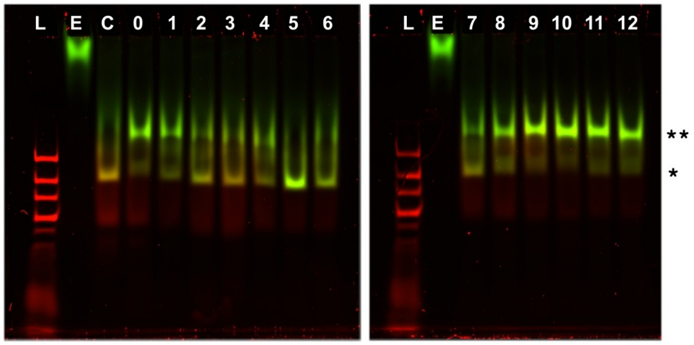


**L**-Ultra Low range Ladder
**E**- H8TEV_EGFP_2xDBP_30AAFL
(abbreviated as Protein)

**C**-Protein + 34-dplxDimerCntrl
**0**-Protein + 34-dplxDimer_0
**1**-Protein + 34-dplxDimer_1
**2**-Protein + 34-dplxDimer_2
**3**-Protein + 34-dplxDimer_3
**4**-Protein + 34-dplxDimer_4
**5**-Protein + 34-dplxDimer_5
**6**-Protein + 34-dplxDimer_6
**7**-Protein + 34-dplxDimer_7
**8**-Protein + 34-dplxDimer_8
**9**-Protein + 34-dplxDimer_9
**10**-Protein + 34-dplxDimer_10
**11**-Protein + 34-dplxDimer_11
**12**-Protein + 34-dplxDimer_12

**c** Heating and cooling protocols

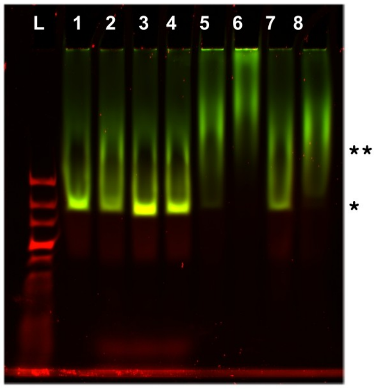


| Lane | Duplex | Heating | Cooling to 23 °C |
|---|---|---|---|
| 1 | 34-5 | 37 °C, 10 min | 0.1 °C steps over 1 h |
| 2 | 34-6 | 37 °C, 10 min | 0.1 °C steps over 1 h |
| 3 | 34-5 | 55 °C, 10 min | 0.1 °C steps over 1 h |
| 4 | 34-6 | 55 °C, 10 min | 0.1 °C steps over 1 h |
| 5 | 34-5 | 37 °C, 10 min | 0.1 °C steps over 12 h |
| 6 | 34-6 | 37 °C, 10 min | 0.1 °C steps over 12 h |
| 7 | 34-5 | 55 °C, 10 min | 0.1 °C steps over 12 h |
| 8 | 34-6 | 55 °C, 10 min | 0.1 °C steps over 12 h |

L, Ultra Low range Ladder. Lanes 1–8, Protein + indicated duplex.

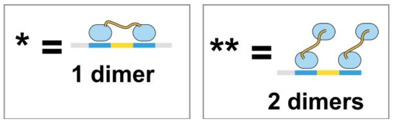


**Supplementary Figure S2. Annealing and cooling of the 30AA *cis*-dimer.** DNA strands were annealed at a 1:1 molar ratio and mixed with fourfold excess H8TEV–EGFP–2×DBP35opt–30AAFL. (a) Room-temperature mixing. (b) Heating to 55 °C for 10 min. (c) Heating temperatures and cooling protocols listed in the table. Samples were analyzed by 6% native PAGE at 100 V for 120 min in an ice bath. Sequences are provided in Supplementary_Tables.xlsx.

**a** **E**- H8TEV_EGFP_2xDBP_51AAFL (abbreviated as Protein)

**b**

**c** **E**- H8TEV_EGFP_2xDBP_72AAFL (abbreviated as Protein)

**d**

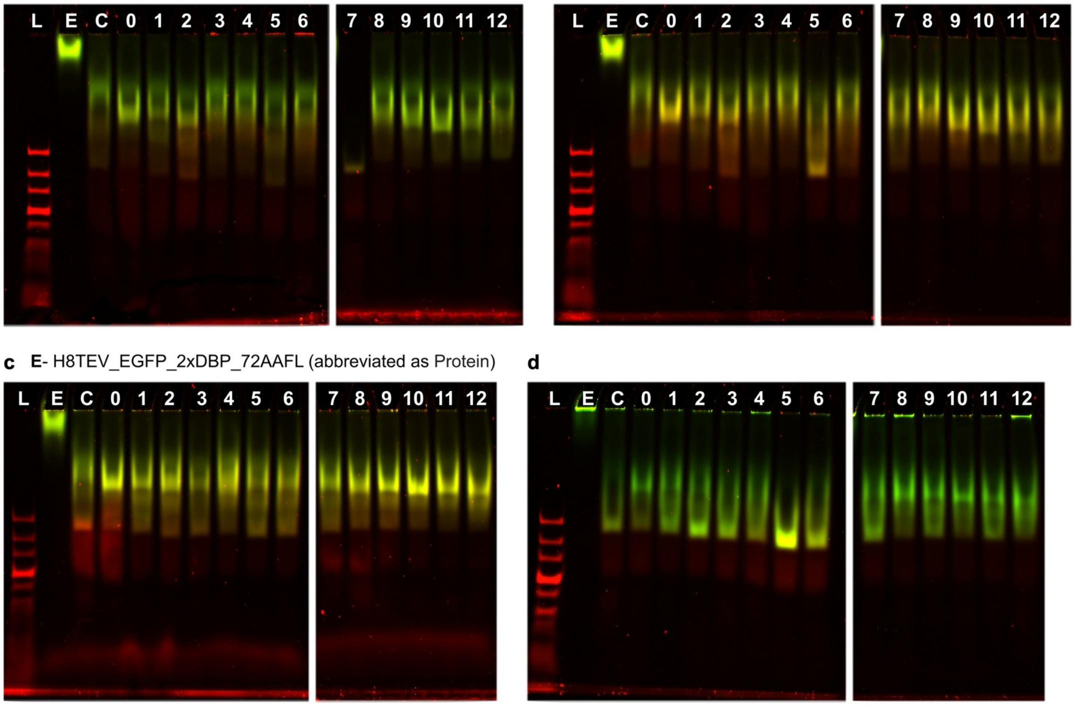


**L**-Ultra Low range Ladder
**C**-Protein + 34-dplxDimerCntrl
**0**-Protein + 34-dplxDimer_0
**1**-Protein + 34-dplxDimer_1
**2**-Protein + 34-dplxDimer_2

**3**-Protein + 34-dplxDimer_3
**4**-Protein + 34-dplxDimer_4
**5**-Protein + 34-dplxDimer_5
**6**-Protein + 34-dplxDimer_6

**7**-Protein + 34-dplxDimer_7
**8**-Protein + 34-dplxDimer_8
**9**-Protein + 34-dplxDimer_9
**10**-Protein + 34-dplxDimer_10

**11**-Protein + 34-dplxDimer_11
**12**-Protein + 34-dplxDimer_12

**e**

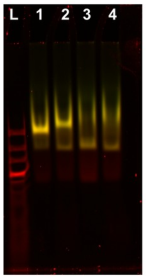


| Lane | Duplex | Treatment |
|---|---|---|
| **1** | 34-2 | RT |
| **2** | 34-3 | RT |
| **3** | 34-2 | 40 °C for 10 min, then RT for 10 min |
| **4** | 34-3 | 40 °C for 10 min, then RT for 10 min |

L, Ultra low range ladder.
Duplexes mixed with 2xDBP35no_linker.

**f**

37°C -10 min and brought down with 0.1°C steps to 23°C, over 1 hour

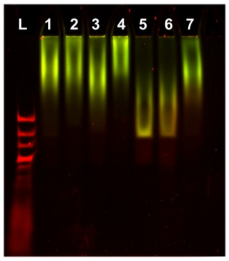


**L**-Ultra Low range Ladder
**1**-6AAHL + 34-5 duplex
**2**-6AAHL + 34-7 duplex
**3**-11AAHL + 34-2 duplex
**4**-11AAHL + 34-7 duplex
**5**-30AAFL + 34-5 duplex
**6**-30AAFL + 34-6 duplex
**7**-51AAFL + 34-5 duplex

**g**

55°C -10 min and brought down with 0.1°C steps to 23°C, over 1 hour

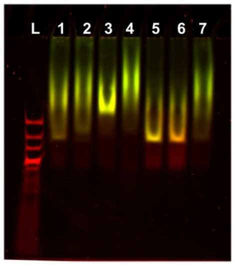


**h**

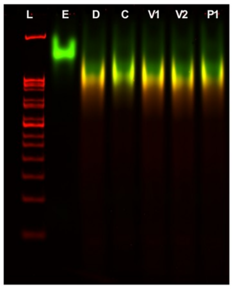


**L**-50bp Ladder
**E**-H8TEV_2xDBP48_30AAFL (abbreviated as Protein)

**D**-Protein + 48-dplx_5
**C**-Protein+ 48-dplx_cntrl
**V1**-Protein + 48-dplx_5V1
**V2**-Protein + 48-dplx_5V2
**P1**-Protein+ 48-dplx_5pali

**Supplementary Figure S3. Additional *cis*-dimer linker constructs.** (a–d) 51AA (a, b) and 72AA (c, d) flexible-linker *cis*-dimers at room temperature (a, c) and after heating to 55 °C for 10 min (b, d). (e) No-linker DBP35opt *cis*-dimer across annealing conditions. (f, g) Selected DBP35opt linker constructs. (h) DBP48 control construct. Samples were analyzed by 6% native PAGE at 100 V for 120 min in an ice bath.

**a** in FOB buffer  **b** in PBS buffer

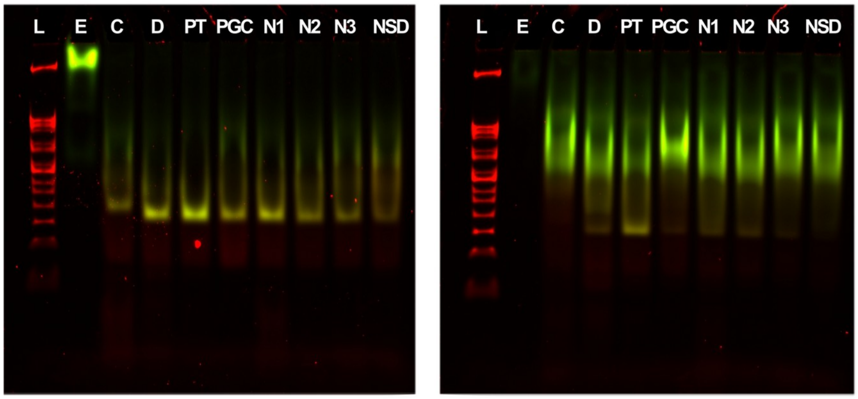


**L**-Ultra Low range Ladder
**E**- H8TEV_EGFP_2xDBP_30AAFL (abbreviated as Protein)

**C**-Protein + 34-dplxDimerCntrl2
**D**-Protein + 34-dplxDimer_5
**PT**-Protein + 34-dplxDimer_PT
**PGC**-Protein + 34-dplxDimer_PGC
**N1**-Protein + 34-dplxDimer_Nick1
**N2**-Protein + 34-dplxDimer_Nick2
**N3**-Protein + 34-dplxDimer_Nick3
**NSD**-Protein + 34-dplxDimer_NickSD

**c**

| | | |
|---|---|---|
| **34.dplxDimer.5f:**<br>**34.dplxDimer.5r:** | GCATATATTGCACATATATTGCACAATATATAGC<br>GCTATATATTGTGCAATATATGTGCAATATATGC | **34-dplxDimer_5** |
| **34.dplxDimer.5f:**<br>**34.Dimer.5rNick1a:**<br>**34.Dimer.5rNick1b:** | GCATATATTGCACATATATTGCACAATATATAGC<br>ATATATGTGCAATATATGC<br>GCTATATATTGTGCA | **34-dplxDimer_Nick1** |
| **34.dplxDimer.5f:**<br>**34.Dimer.5rNick2a:**<br>**34.Dimer.5rNick2b:** | GCATATATTGCACATATATTGCACAATATATAGC<br>ATATGTGCAATATATGC<br>GCTATATATTGTGCAAT | **34-dplxDimer_Nick2** |
| **34.dplxDimer.5f:**<br>**34.Dimer.5rNick3a:**<br>**34.Dimer.5rNick3b:** | GCATATATTGCACATATATTGCACAATATATAGC<br>CAATATATGTGCAATATATGC<br>GCTATATATTGTG | **34-dplxDimer_Nick3** |
| **34.Dimer.5f_SD:**<br>**34.Dimer.5rNick1a_SD:**<br>**34.Dimer.5rNick1b_SD:** | GCATATATTGCACATATATTGCACAATACCTAGC<br>ATATGTGCAATATATGC<br>GCTAGGTATTGTGCAAT | **34-dplxDimer_NickSD** |

**d**

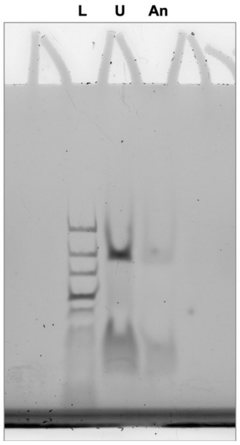


| Lane | Sample | Treatment before loading | Loaded |
|---|---|---|---|
| **L** | Ladder | — | — |
| **U** | 34-duplexDimer_5 + fourfold excess H8TEV–2xDBP–30AAFL | Room temperature, 10 min; not heated | 400 nM |
| **An** | Same mixture as U | 55 °C, 10 min; band excised from gel and purified; 3 kDa MWCO spin at 5,000 rcf | Nominal 400 nM, from DNA absorbance; signal weaker than U |

**Supplementary Figure S4. Effects of nicking, flanking sequence and gel extraction on 30AA *cis*-dimer binding.** (a, b) Binding to nicked and flanking-sequence duplexes in FOB (a) and PBS (b); lane abbreviations are listed to the right of the gels. DNA strands were annealed at 1:1 or 1:1:1 molar ratios, mixed with fourfold excess protein, heated to 55 °C for 10 min and cooled to room temperature for 10 min. (c) Reference and nicked duplexes; recognition sites are blue and spacers are gold. (d) Gel extraction and re-electrophoresis of 34-duplexDimer_5 mixed with fourfold excess H8TEV–2×DBP35opt–30AAFL (L, ladder; U, unheated mixture; An, gel-extracted annealed band). Gels in a and b were 6% native PAGE (100 V, 120 min, ice bath). All sequences are provided in Supplementary_Tables.xlsx.

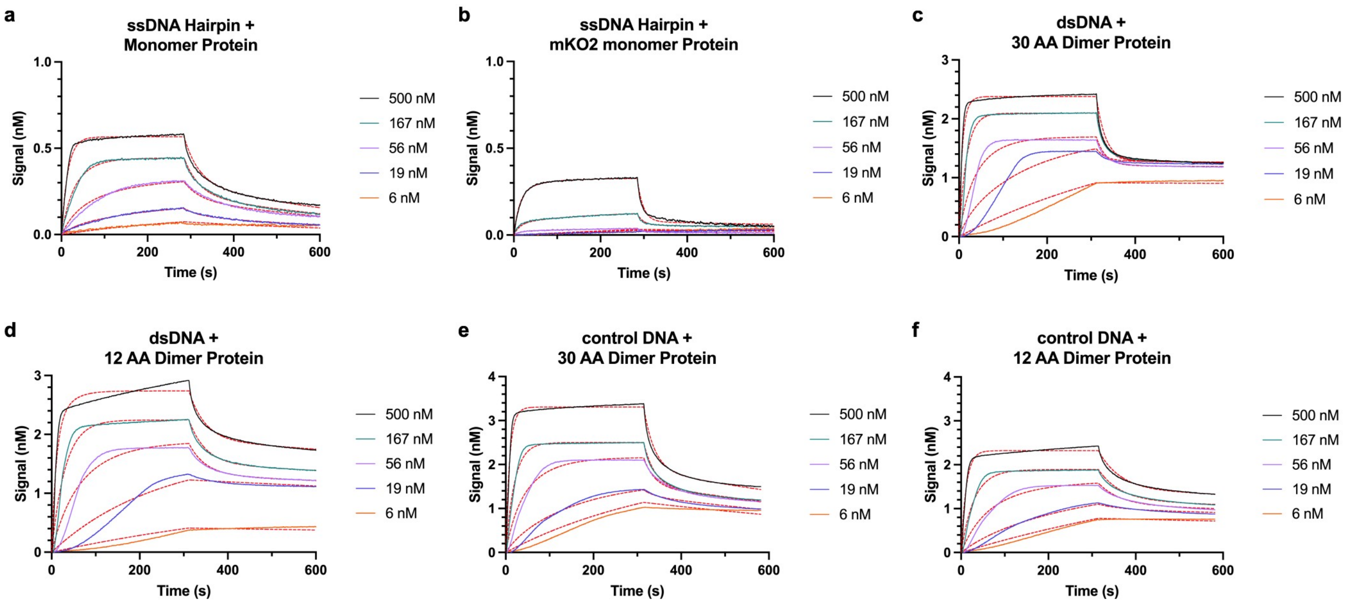


**Supplementary Figure S5. Biolayer interferometry.** (a) Monomer with hairpin DNA. (b) mKO2 monomer fusion with hairpin DNA. (c, d) 30AA (c) and 12AA (d) *cis*-dimers with target duplex DNA. (e, f) 30AA (e) and 12AA (f) *cis*-dimers with control DNA. Solid curves show measured responses and red dashed curves show fitted curves; protein concentrations are indicated. Kinetic parameters and DNA sequences are provided in Supplementary Table S1 and Supplementary_Tables.xlsx.

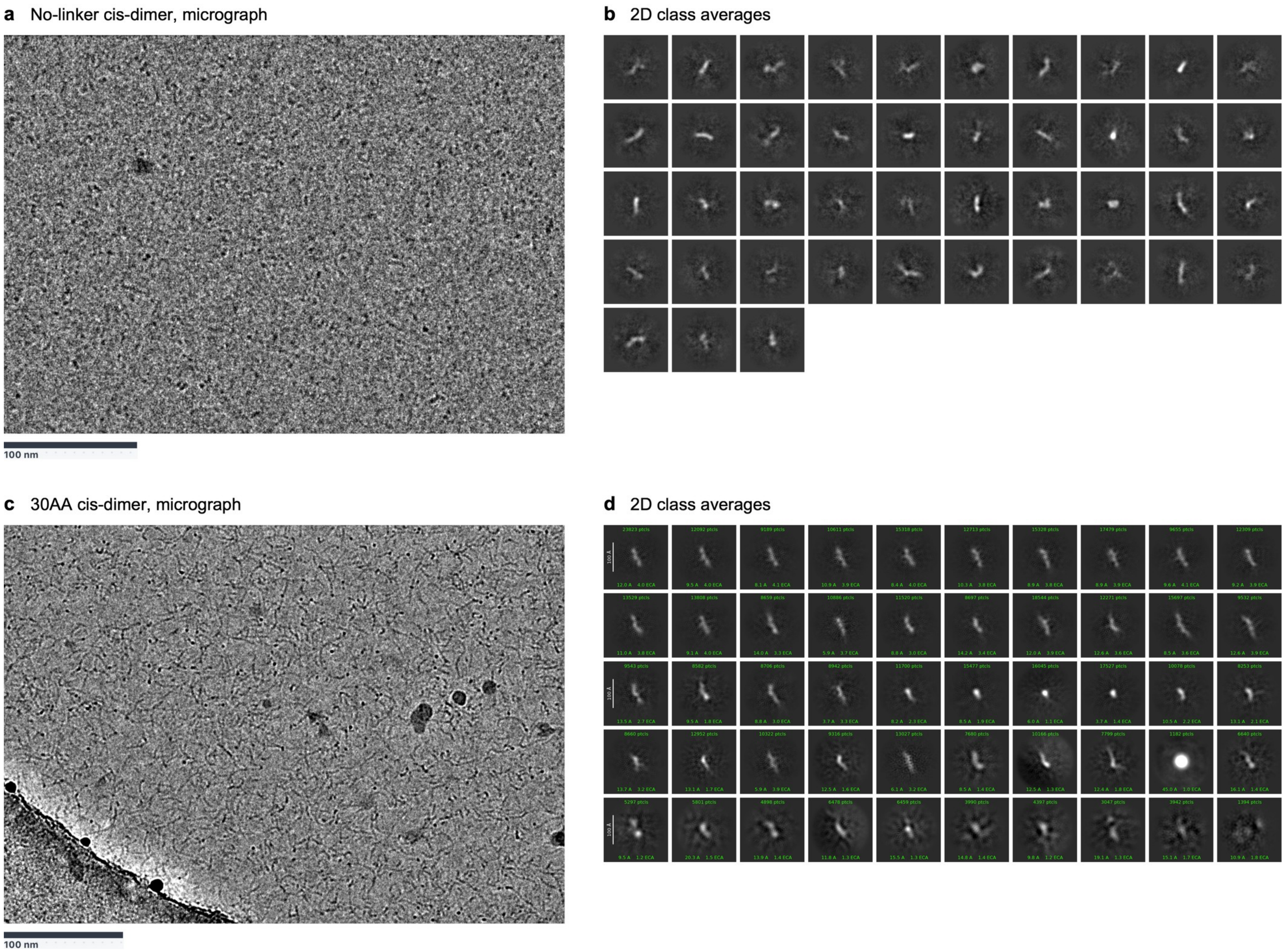


**Supplementary Figure S6. Cryo-EM micrographs and 2D class averages of *cis*-dimer complexes.** (a, b) 34-2 duplex with H8TEV–EGFP–2×DBP35opt without a linker (1:1 DNA:protein ratio, 3 µM DNA): representative micrograph (a) and selected 2D class averages (b). (c, d) 34-5 duplex with H8TEV–EGFP–2×DBP35opt–30AAFL (1:0.5 DNA:protein ratio, 3 µM DNA): representative micrograph (c) and selected 2D class averages (d). A high-resolution 3D reconstruction was not obtained from either dataset, likely due to the small size and conformational flexibility of the complexes. Scale bars, 100 nm.

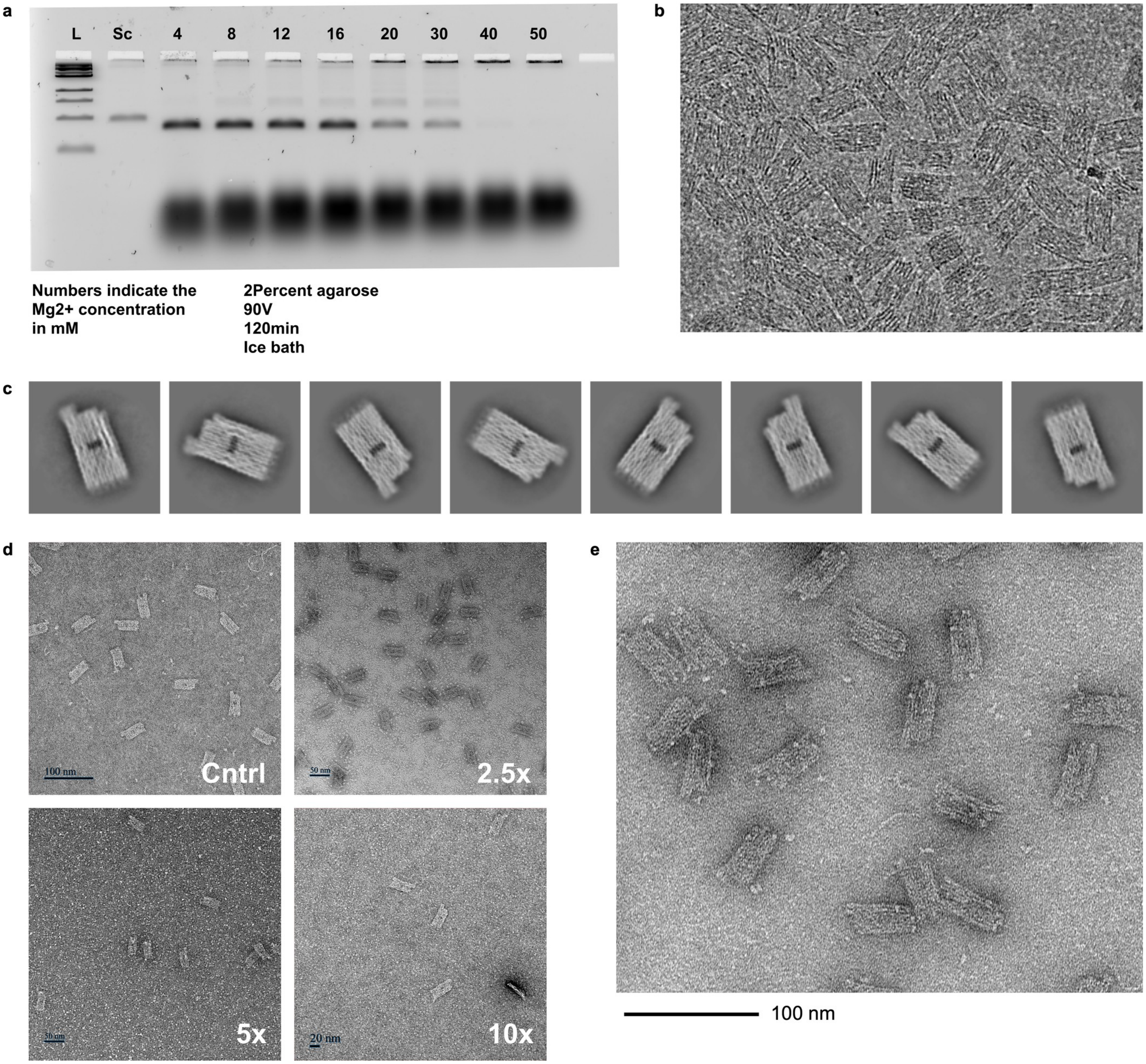


**Supplementary Figure S7. Q1 origami characterization and origami-bound particles.** (a) Magnesium screen for Q1 origami folding (2% agarose gel). (b) Cryo-EM micrograph of Q1 incubated with a fivefold molar excess of H8TEV–EGFP–DBP35opt. (c) Cryo-EM 2D class averages showing no reliable density of bound monomeric protein. (d) Negative-stain TEM of Q1 with increasing concentrations of H8TEV–EGFP–DBP35opt; scale bars are shown in the images. (e) Negative-stain TEM overview accompanying the Q1+D4 cryo-EM class average in Fig. 5d; scale bar, 100 nm.

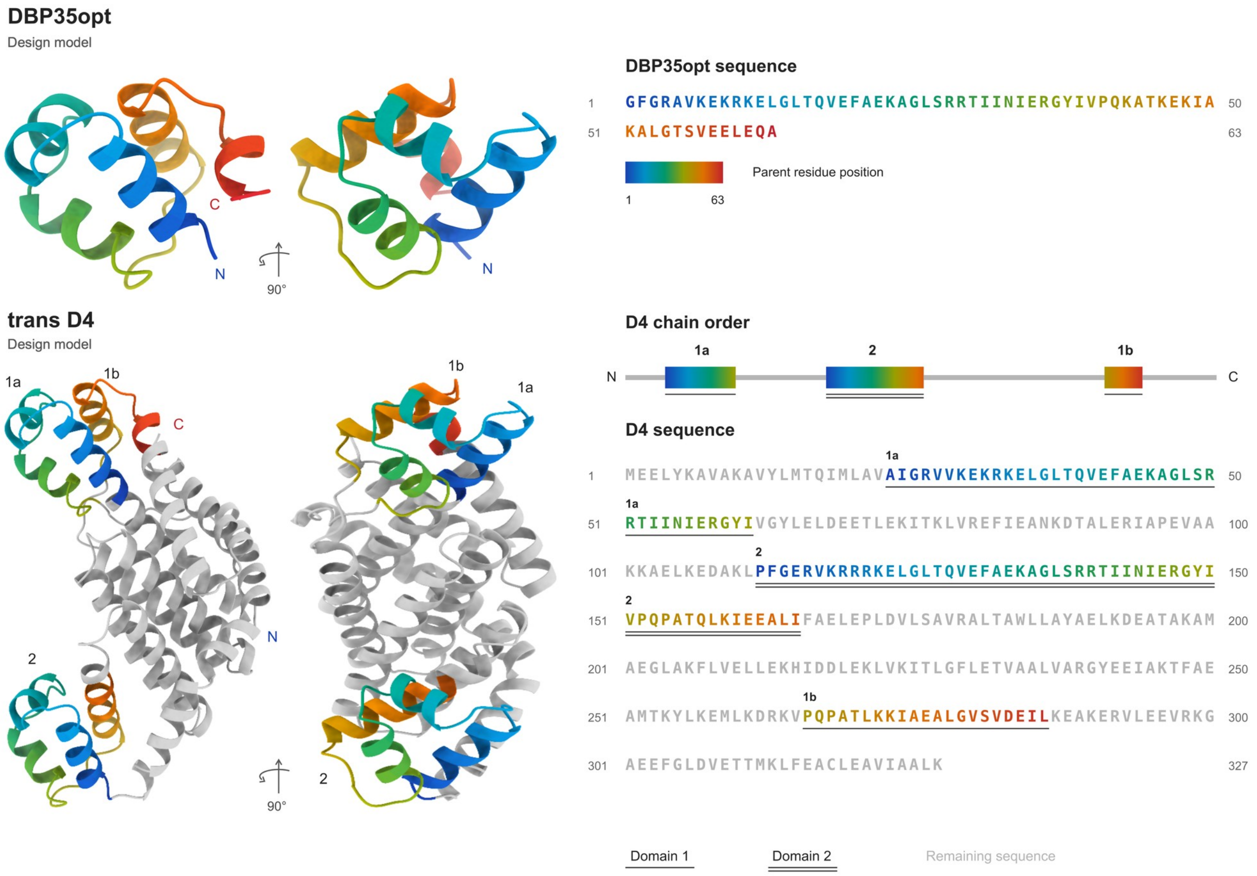


**Supplementary Figure S8. DBP35opt and D4 *trans*-dimer design models and sequence correspondence.** Top, two views (90° rotation) of the 63-residue parent DBP35opt design model colored from N terminus (blue) to C terminus (red) alongside its amino-acid sequence. Bottom, two views (90° rotation) of the 327-residue circularly permuted D4 *trans*-dimer design model, chain-order diagram showing split domain 1 (1a, 1b) flanking domain 2 (2), and full D4 sequence. Rainbow colors indicate residues corresponding to parent DBP35opt positions; gray denotes *de novo* designed bridge and terminal-extension residues.

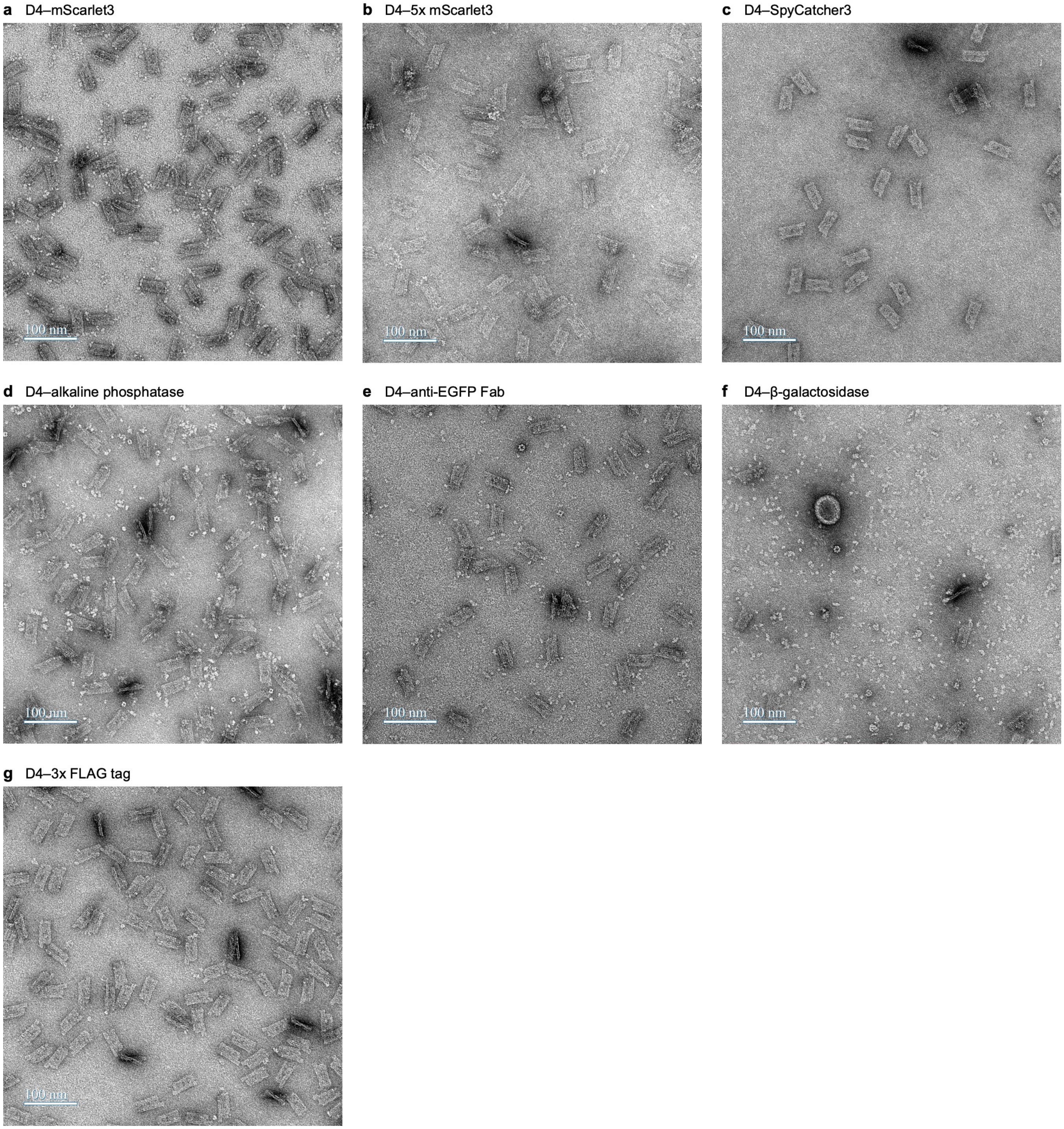


**Supplementary Figure S9. Negative-stain TEM of D4–cargo fusions on Q1 origami.** Representative negative-stain TEM fields of Q1 with D4–mScarlet3 (a, 24-fold excess), D4–5× mScarlet3 (b, 24-fold), D4–SpyCatcher3 (c, sixfold), D4–alkaline phosphatase (d, 24-fold), D4–anti-EGFP Fab (e, 48-fold), D4–β-galactosidase (f, 24-fold) and D4–3× FLAG tag (g, 24-fold). Samples contained 10 nM Q1 in 5 µL and were incubated at 4 °C for 1 h. Scale bars, 100 nm. Full gel series and additional micrographs are deposited in Zenodo (https://doi.org/10.5281/zenodo.24519036).

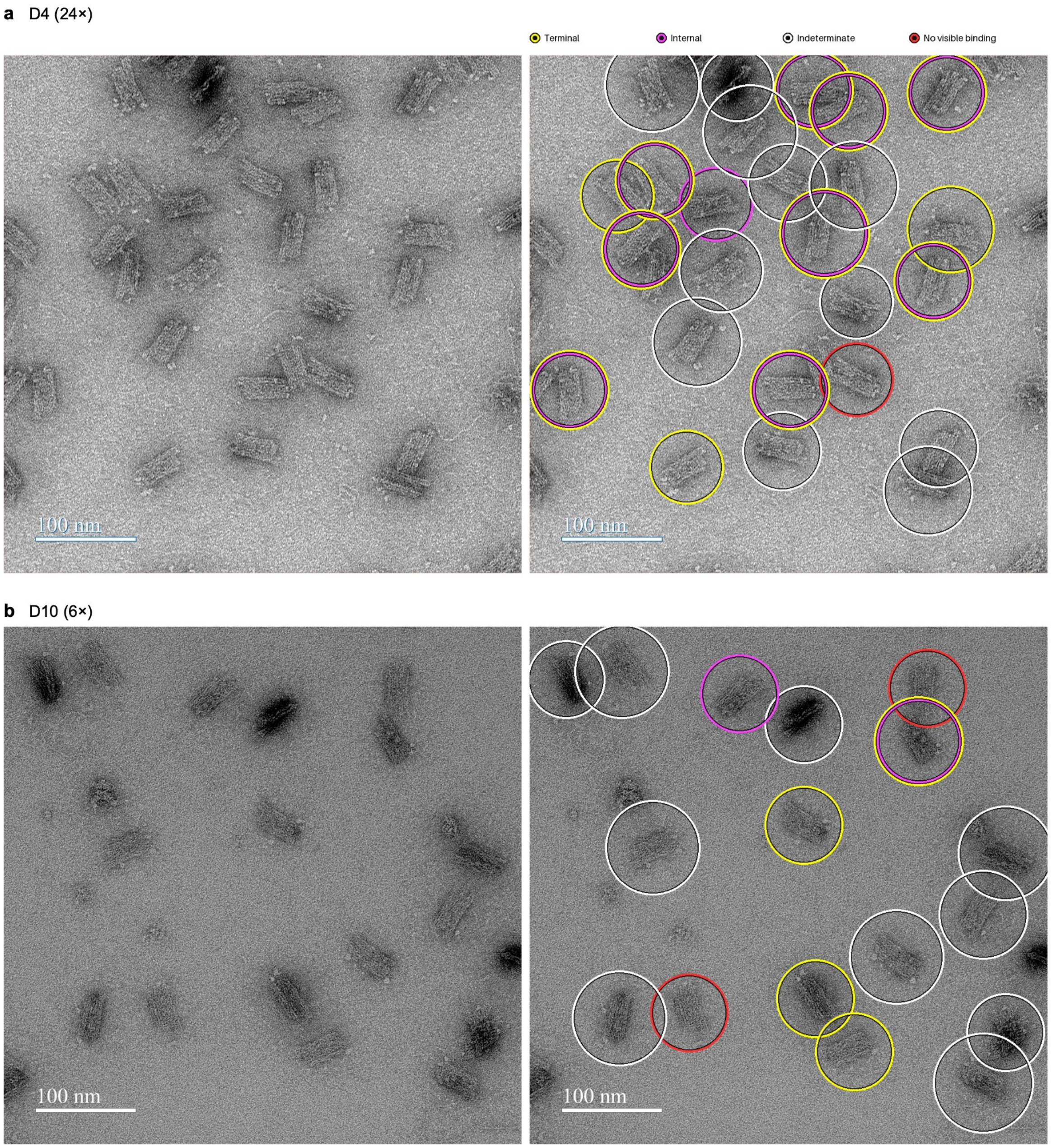


**Supplementary Figure S10. Trans-dimer occupancy screen on Q1 origami.** Negative-stain TEM of Q1 with D4 at 24-fold excess (a) and D10 at sixfold excess (b). Each field is shown without overlays on the left and with simplified manual annotations on the right (yellow, terminal; magenta, internal; white, indeterminate; red, no visible binding; terminal and internal labels may coexist). Scale bars, 100 nm. Complete micrographs and manual annotations for all four screened designs (D4, D34, D10 and D46) are provided in Supplementary Data 1 (Zenodo, https://doi.org/10.5281/zenodo.24519036).
